# The use of pigs vocalisation structure to assess the quality of human-pig relationship

**DOI:** 10.1101/2022.03.15.484457

**Authors:** Avelyne S Villain, Carole Guérin, Céline Tallet

**Affiliations:** PEGASE, INRAE, Institut Agro, 35590 Saint Gilles, France; Behavioural Ecology Group, Section for Ecology & Evolution, Department of Biology, University of Copenhagen, 2100, Copenhagen Ø, Denmark

**Keywords:** Key words, Positive handling, Acoustic communication, Emotions, Mood, Behaviour, Welfare, Interspecific interactions.

## Abstract

Studying human-animal interactions in domestic species and how they affect the establishment of a positive Human-Animal Relationship (HAR) may help us improve animal welfare and better understand the evolution of interspecific interactions associated with the domestication process. Understanding and describing the quality of an HAR requires information on several aspects of the animal biology and emotional states (social, spatial and postural behaviours, physiological and cognitive states). Growing evidence shows that acoustic features of animal vocalisations may be indicators of emotional states. Here, we tested the hypothesis that vocal structure may indicate the quality of HAR. At weaning, 30 piglets were positively handled by an experimenter who talked to and physically interacted with them three times a day, while 30 other piglets only received the contact necessary for proper husbandry. After two weeks, we recorded the behaviours and vocalisations produced in the presence of the static experimenter for 5 min. We repeated this test two weeks later, after a conditioning period during which human presence with additional positive contacts was used as a reward for all piglets. We hypothesized this conditioning period would lead to a positive human-piglet relationship for all piglets. As expected, piglets that were positively handled at weaning expressed a higher attraction toward the experimenter, and, after the conditioning, piglets that were not positively handled at weaning expressed a similar level of attraction than the positively handled ones. Piglets positively handled at weaning produced shorter grunts than the other ones, regardless of the context of recording, which may indicate a more positive affect. During reunions with the static experimenter, a more positive HAR was associated with a decrease in vocal reactivity to human proximity. However, during reunions with the experimenter providing additional positive contacts and over the conditioning, proximity to the human systematically triggered shorter and higher pitched grunts, indicator of positive a emotional state. Results first show that changes in vocal structure are consistent with indicators of positive states in the presence of a human. Second, these changes are stronger when the human positively interact with the piglets, supposedly emphasizing a higher positive arousal state during these interactions. We show that vocalisation structure may be a promising indicator of the quality of human-pig relationship.

## Introduction

The process of domestication was conducted to shape physiology and morphology of domestic animal species, but also their behaviour. It notably has shaped interspecific interactions between human and non-human animals, by improving animals’ capacity to use human signals to adapt their behaviour both decreasing fearfulness toward humans and increasing attention toward humans (Mignon-Grasteau et al., 2005). In farms, the relationship that domestic animals form with humans is important for animal welfare. Therefore, studying human-animal interactions and their consequences to understand the mechanisms of emergence and maintenance of a positive human-animal relationship (HAR) directly applies to welfare (Rault et al., 2020). Animal welfare consists of three major aspects: the ability of an animal to control its mental and physiological stability (Broom, 2011), the decrease of experiencing negatively perceived contexts and the increase in experiencing positively perceived contexts and species-specific behaviors (Peterson et al., 1995; Weerd & Day, 2009). A positive HAR is thought to be established through repeated positive interactions between the human and the non-human animal. Some of the mechanisms involved in this process are: accumulation of positive experiences through positive associative learning, modifications of cognitive biases, shaping expectations from the non-human animal toward the human. A positive HAR can be appreciated through behavioural and physiological measures, for example by assessing the expression of positive emotions [reviewed in (Rault et al. 2020)]. Several behavioural measures may help to define a positive HAR such as: short latency to approach and spatial proximity (Boivin et al., 2000; Schmied et al., 2008), body postures (Villain, Lanthony, et al., 2020) or play behaviour (Jerolmack, 2009). Contacts from a human such as stroking, may induce changes in body postures and exposition of body areas by the animal to the human, supposedly vulnerable [central neck area in cattle (Schmied et al. 2008), abdominal area in pigs (Rault et al., 2019)]. Such grooming solicitation may be markers of engagement, trust and motivation to interact with the human. In most cases, these behaviours are similar to those shown during intraspecific socio positive interactions, although there are some species specific behaviours [e.g., dog vs. wolf (Gácsi et al., 2005)]. Vocal behaviour may also help defining the quality of an HAR. First, some vocalisations type have been associated with positive interactions with humans, for example the cat – human communication : purring is thought to be derived from mother pup communication during nursing and is observed associated with care solicitation from humans; meowing, which is not observed during intra specific interactions is thought to emerge from associative learning during cat – human interactions (Brown & Bradshaw, 2014). This shows that HAR may elicit specific vocalisations from the non human animal toward the human. Second, vocalisation structure is known to carry markers of the emotional states in several bird and mammal species (Briefer, 2012, 2020) and markers of emotional valence (positive versus negative) has been studied in domestic farm animals [reviewed in Laurijs et al. (2021)]. Since positive or negative HAR is likely to affect the emotional state of animals, it is likely that it may be reflected in the structure of the produced vocalisations.

In pigs, diversified evidence attest the possibility of a positive HAR. Animals may be handled by humans providing regular additional positive contacts, leading to the expression of a positive perception of humans, with evidence from behavioural and physiological studies. Cognitive bias tests showed a positive judgment bias in piglets that had received gentle contacts with humans (Brajon et al., 2015b). Pigs may recognise a human providing positive contacts compared to an unfamiliar one and adapt their behaviour accordingly (Brajon et al., 2015c). Pigs may be sensitive to human voice and respond accordingly (Bensoussan et al., 2019, 2020). Pigs vocalisations are diverse and linked to their emotional states, attested by the use of positive or negative call types (Briefer et al., 2019, 2022; Tallet et al., 2013). In addition, even within a call type, spectro-temporal changes are closely related to the valence or the arousal a situation may trigger for the animal. For example, the grunt, a contact call, is used in various contexts and is now known to be a flexible call. Positive situations have been associated with shorter grunts compared to negative ones (Briefer et al., 2019, 2022; Friel et al., 2019), as well as higher formants (which are frequency peaks containing more energy than others) and a lower fundamental frequency during positive situations (Briefer et al., 2019, 2022). Grunt structure may also change according to the arousal of a negative situation: the higher the arousal in the negative state the higher the frequency range and bandwidth (Linhart et al., 2015) and the longer (Puppe et al., 2005) the grunts. Variation in grunt spectro-temporal structure in positive situations of different arousal is still unknown.

In order to determine to what extent vocalisations structure could be used as non invasive indicator of the quality of human-pig relationship, we tested whether varying the degree of familiarity and the quality of the human-pig interactions could modulate the spectro-temporal structure of vocalisation, through the vocal expression of emotional state. Because it was suggested to study vocal markers of emotions within the same call type (Briefer, 2020) and because grunts are the most commonly produced call in various contexts, we studied the spectro-temporal structure of grunts. We predicted that if grunts reflect the quality of the human-pig relationship, then 1. A period of positive handling given by a human should modulate piglets vocal expression in presence of the human, leading to grunts exhibiting markers of positive states (higher pitched and shorter grunts), 2. Spatial proximity toward the human should influence the spectro-temporal structure of grunts (higher pitched and shorter grunts).

## Methods

### Ethical note

The study was approved by the ethic committee CREEA and received the authorization no. APAFIS#17071-2018101016045373_V3 from the French Ministry of Higher Education, Research and Innovation. UE3P, where the experiment was carried out, is an experimental unit authorized by the French Ministry of Agriculture to breed animals for experimentation under the number D35-275-32. This authorization includes a derogation to follow the directive 2008/120/EC relative to the protection of piglets and its regulations.

### Subjects and housing conditions

Sixty weaned female pigs (in two replicates from January to April 2019), *Sus scrofa domesticus*, bred from crosses between Large White and Landrace females and Piétrain males were used for this study from 28 to 62 days after birth. Animal housing and experiments took place at the experimental unit UE3P (UE 1421, INRAE France).

One piglet had to be excluded from our sample size to receive care/medication due to health issues independent from the experiment. From weaning at 28 days of age, piglets from the same litter and having similar weight (<1 kg difference) were housed by three in a 1.2 x 1.3m pen on plastic duckboard. Wooden panels were used to visually isolate pens. One metal chain per pen was used for enrichment. Food and water were available *ad libitum*. Artificial lights were turned on from 8:00 to 17:00 and temperature was maintained between 26 and 27 °C. The experiment was carried out in two replicates and two identical rearing rooms were used (5 pens per room per replicate).

### Treatment: positive handling at weaning

From day 28 (day of weaning) to day 39 of life, piglets were separated into two groups that experienced a different post-weaning period as follows:

#### - Non positively handled piglets (H piglets)

- Control piglets from 10 rearing pens, housed in the same room, received the minimal amount of daily contact with a stockperson (a 1.70m tall male who did the feeding, cleaning and health checkups). The stockperson wore a dark green shirt and pants and brown shoes.
- Positively handled piglets piglets (H+ piglets):
- Experimental piglets from the 10 other rearing pens, housed in another room, received the same daily care given by the same stockperson as for H piglets. They additionally received repeated sessions of additional human contacts. Each pen of three piglets received 29 sessions of 10 min, from day 28 (weaning) until day 39, occurring five days a week. Three sessions per day were performed (except on the day of weaning during which only two were done with a two-hour break in between). Each session took place in the rearing pen and the order of the interventions in the pens was balanced across days. The handling procedure, using gentle tactile contacts is described in supplementary material of Villain et al. (2020) and was similar to Tallet et al. (2014). Briefly, the behaviour of the human toward the piglet was adapted to the reaction of each animal and included four steps: (1), the handler hold out the hand towards the animal; (2) if the piglet did not move away, the handler tried to touch it; (3) if the piglet accepted being touched, the handler softly stroked it along the body with the palm of her hand; and (4) once it accepted being stroked, the handler scratched it along the body with her fingers. Scratching consisted in rubbing the skin of the piglets with the finger tips and applying more pressure than stroking. No specific body part of the piglets was more considered that another. Two experimenters (‘AV’ and ‘AH’) performed these sessions (both women, both between 1.70-1.73 m tall, with a balanced number of pens attributed to each of them). The experimenters wore the same blue overalls and green boots each time they interacted with the piglets. The experimenters tried to imitate each others behaviours (remote video monitoring) to decrease variability.

This intense period of additional positive contacts for half of the piglets after weaning constituted the treatment of positive handling at weaning: positively handled piglets are referred to as H+ piglets and non positively handled piglets are referred to as H piglets to describe the early experimental treatment they experienced regarding a human, prior to the conditioning.

#### Conditioning: sessions of additional positive contacts with (un)familiar human

The conditioning took place between day 42 and 62 of age and lasted twelve days, with two trials per day and at least three hours between trials on the same day. Piglets were habituated to the test room for 10 min, by pen, two days before the start of the conditioning. All piglets (H and H+) were subjected to the same conditioning. The experimental design of the conditioning is already published in an article dedicated to the study of anticipatory behaviour (Villain, Hazard, et al., 2020).

Briefly, all piglets were individually trained to learn to associate two different stimuli with the arrival of two different (pseudo)-social partners: either two pen mates (partner = Conspecifics) or a familiar human (partner = Human). When entering the room, the piglets and the partner(s) would remain in the room for 2 min. Specifically, when the human was the partner, the human entered, sat on a bucket and positively interacted with the piglet for 2 min, in the same manner as additional contacts was provided to the H+ piglets during the previous period (see above section) (figure 1). Therefore, at the beginning of the conditioning, H+ piglets were already familiar with the human and procedure, whereas H piglets were unfamiliar with the human. During the conditioning, the same sessions occurred in both treatment groups (H and H+). After the conditioning, all piglets were familiar with the human, but treatment groups had a different time of exposure to them. Sessions of reunions with social partners were not studied here because they were part of an analysis on vocal expression of positive anticipation reported earlier (Villain, Hazard, et al., 2020).

**Figure 1:**
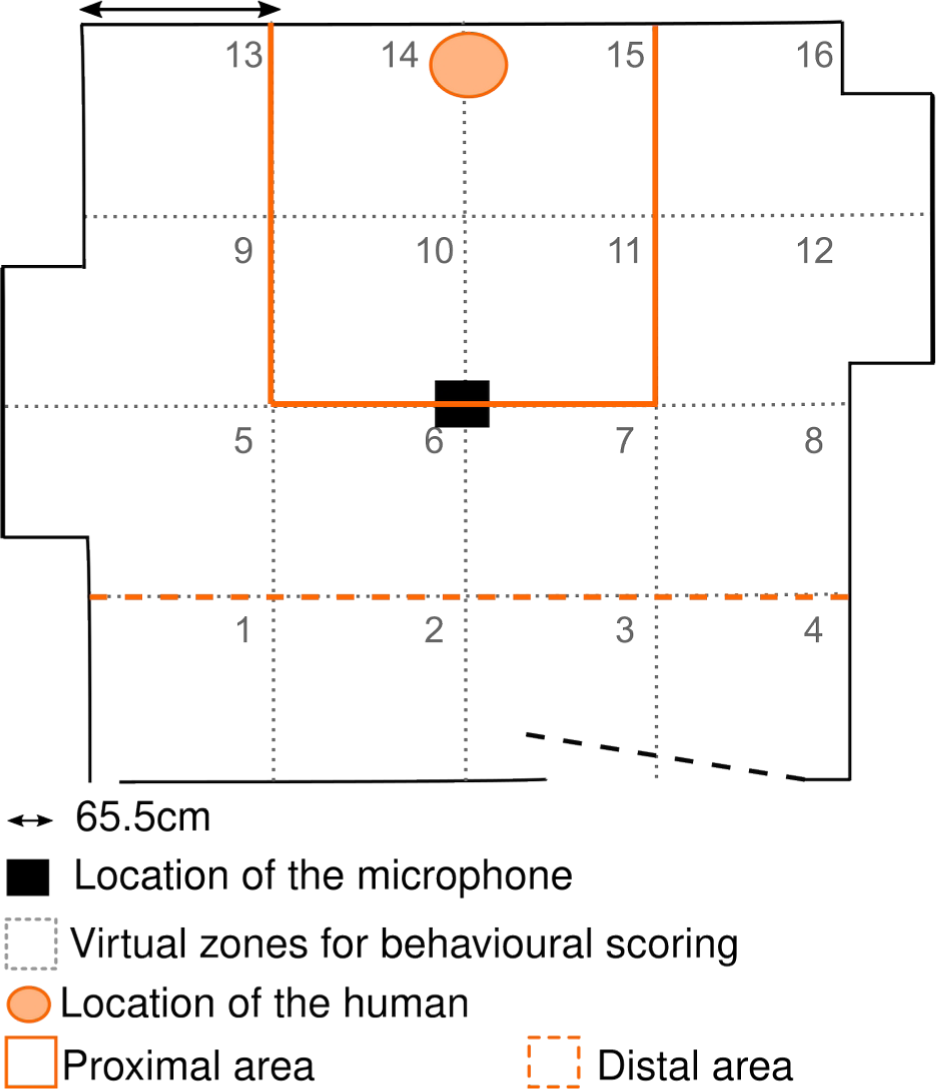
Design of the room used during the Isolation/Reunion tests and the additional positive contacts sessions of the conditioning. The room was split into 16 virtual zones. A proximal area (zones 10, 11, 14, 15) and a distal area (zones 1, 2, 3, 4) were defined, suing the location of the human as reference. Behavioural monitoring and analyses

For every second trial, the 2 min reunions with the human were analysed by the same person: trials number 2, 4, 6, 8, 10 and 11 (see behavioural analyses section).

#### Standard Isolation/Reunion Tests with a static and silent human

At 40 or 41 (before conditioning) and then 63 or 64 (after conditioning) days of age, piglets were subjected to a standard Isolation/Reunion test in order to assess their perception of the human. The test consisted of two phases. The piglet was brought individually in a trolley to the experimental room. It was left alone for 5 min, which defined the ‘Isolation’ phase. Then, the human entered the room, remained stand up for 30 seconds and they sat on a bucket, remaining silent and not moving for 4.5 min (figure 1).

Sessions and tests were recorded using a camera (Bosh, Box 960H-CDD) and behaviours were scored *a posteriori* on videos using *The Observer XT 14.0* (Noldus, The Netherlands) software. The room was split into 16 virtual equally-dimensioned zones to assess the mobility and exploratory behaviour of the piglet. A proximal area, around the human was defined by merging four zones, a distal area was defined merging the four most distant zones from the human (figure 1).

The behaviours scored during the reunion of the Isolation/Reunion test and the sessions of additional positive contacts of the conditioning are available in table 1. Every time the shoulders of the piglet crossed a zone, a zone change was scored. Looks and watching behaviours were scored as point events, all other behaviours were scored as state events. Behavioural scores were then calculated to quantify global responses (see Table 1).

**Table 1:** Ethogram.

| Behaviour | Description |
| --- | --- |
| No. zones crossed <sup>1,2</sup> | The number of times the piglet crossed a virtual zone |
| No. approaches H <sup>1</sup> | Number of times the piglets entered the proximal area |
| Time watching H <sup>1,2</sup> | The amount of time the piglet spent watching the human |
| Latency to contact H <sup>1,2</sup> | The latency to the first contact of the human by the piglet |
| No. looks toward H <sup>1,2</sup> | The number of times the piglet turned its head toward the human |
| No. looks at walls and doors <sup>1</sup> | The number of times the piglet looked at walls and doors |
| Time watching walls and doors <sup>1</sup> | The amount of times the piglet watched walls and doors |
| Time in proximal area <sup>1,2</sup> | The amount of time the piglet spent in the proximal area |
| Time in distal area <sup>1,2</sup> | The amount of time the piglet spent in the distal area |
| Time in contact H <sup>1,2</sup> | The amount of time the piglet investigated the human |
| Time investigating floor <sup>1,2</sup> | The amount of time the piglet investigated the floor |
| No. contacts H <sup>2</sup> | Number of times the piglet was in contact with the human (initiated by the piglet or the human) |

<sup>1</sup>: Scored during reunions of Isolation/Reunion tests. <sup>2</sup>: Scored during reunions of conditioning sessions

#### Acoustic monitoring and analyses

Vocalisations were recorded with an AKG C314 microphone placed in the center of the room and one meter above the ground, connected to a Marantz MD661MK2 recorder. Vocalisations produced during each phase of the trial were manually annotated according to vocal type (grunt, squeal, bark, scream and mixed calls (Kiley, 1972)), after visual inspection of spectrograms using the ‘Annotate’ function of the Praat software (Boersma & Paul, 2001), version 6.0 from http://www.praat.org/. Checking the occurence of each call type in the several contexts of the study, we confirmed that ‘grunt’ was the call type used in all contexts and by most of the piglets in each context. So only the spectro-temporal structure of grunts was further analysed. For information, a table of the number of each call types recorded in each context as well as the number of individuals involved in the count is presented in the electronic supplementary material. We could not conduct a robust statistical analysis on call type utterance, due to the rarity (per subject and tests) of other vocalisations than grunt. (table S5).

A spectro-temporal analysis was performed with custom-written codes using the Seewave R package (Sueur et al., 2008) implemented in R (R Core Team, 2022). We first studied the spectral properties of the remaining background noise of the experimental room (electric noises and remaining low frequency noises from the rest of the building), using 20 examples of 0.5 second fragments and compared it with the general frequency range of the grunts. To avoid measuring masking effect of the background noise, grunts were filtered using a 0.2-8 kHz bandpass filtering (‘fir’ function). As a consequence, all results presented in this study are on a 0.2-8 kHz frequency range, and no conclusions on frequency components of grunts below this 200 Hz threshold are drawn here. Several acoustic parameters were then extracted from each grunt. To measure grunt duration, a 5% to maximal amplitude threshold was used (‘timer’ function). After normalisation, the following spectral parameters were calculated using the ‘specprop’ function (FFT with Hamming window, window length = 512, overlap = 50%): mean (Q50), first (Q25) and third (Q75) quartiles, interquartile range (IQR), centroid and standard deviation (all in Hz). The grunt dominant frequency (in kHz) was also calculated (‘dfreq’, 50% overlapping FFTs, window length = 512), which is the mean over the grunt duration of the frequencies of highest energy of each window. Frequency peaks were detected and the minimal and maximal peaks were kept as descriptors (‘fpeaks’ function, window length = 512, peak detection threshold = 10% of the normalized amplitude). Measures of noisiness and entropy of the grunts were assessed using: Shannon entropy (sh), Spectral Flatness (Wiener entropy, sfm) and Entropy (H) [combining both Shannon and Temporal envelop entropy, length = 512, Hilbert envelop). Two vocal scores were used: the logarithm of grunt duration and a built-in spectral vocal score with all spectral parameters (see below). A table describing mean and range of variation of each acoustic parameter in the relevant contexts of the study is available in the supplementary material (table S4).

#### Statistical analyses

##### Behavioural and vocal response scores

All measures extracted from videos or sound analysis are named parameters throughout the text. The symmetrical distribution of parameters (behavioural on the one hand and acoustic on the other hand) was visually inspected, and linear transformations were computed when necessary to reach symmetrical distribution (see tables 2, 3, 4). When this criteria was reached, Principal Component Analyses (PCA, one for the behavioural analysis and one for the spectral acoustic analysis) were performed using several parameters to build scores [‘dudi.pca’ function from ‘ade4’ R package (Dray & Dufour, 2007) and ‘inertia.dudi’ function to extract the loadings]. These scores were then used as statistical variables. Indeed, PCAs are generally used to reduce the number of variables included in statistical models. It also generates quantifiable global descriptors of behaviours or acoustic structure, since correlated parameters usually load on the same PC (McGregor, 1992). All PCs having an eigenvalue above one were kept and constituted response scores of behavioural (‘ReuPCs’ and ‘CondPCs’ in table 2 and 3 respectively) and vocal (‘VocPCs’, table 4) parameters. Only the duration of grunts was kept separated from the spectral parameters to keep it as a temporal parameter.

**Table 2:** Percentage of explained variance and relative loadings of parameters on PCs, following the Principal Component Analysis computed on the behaviours scored during the reunion of the Isolation/Reunion test. The first three PCs, having an eigenvalue above 1, constituted three behavioural scores: ReuPC1, ReuPC2, ReuPC3. Parameters that explain the most each PC are bolded (|loading|>0.4).

|  | ReuPC1 | ReuPC2 | ReuPC3 |
| --- | --- | --- | --- |
| Cumulative variance explained % | 38.3 | 60.8 | 74 |
| No. of zones crossed (sqrt) | 24.177 | **-55.843** | -0.435 |
| No. approaches H (ln) | **47.748** | -30.163 | 0.578 |
| Time watching H (ln) | **-52.914** | -7.422 | 25.585 |
| Latency to contact H (ln) | **-64.232** | -0.464 | 1.688 |
| No. looks toward H (sqrt) | -7.787 | **-43.721** | 31.633 |
| Time watching room | -32.048 | -13.581 | -6.238 |
| No. looks at walls and doors (sqrt) | 3.524 | **-72.408** | -2.027 |
| Time in proximal area (sqrt) | **69.96** | -0.156 | 9.584 |
| Time in distal area (sqrt) | **-46.416** | -12.437 | -1.215 |
| Time in contact H (sqrt) | **61.041** | 3.586 | 24.183 |
| Time spent investigating floor | 11.868 | -7.503 | **-42.265** |

**Table 3:**
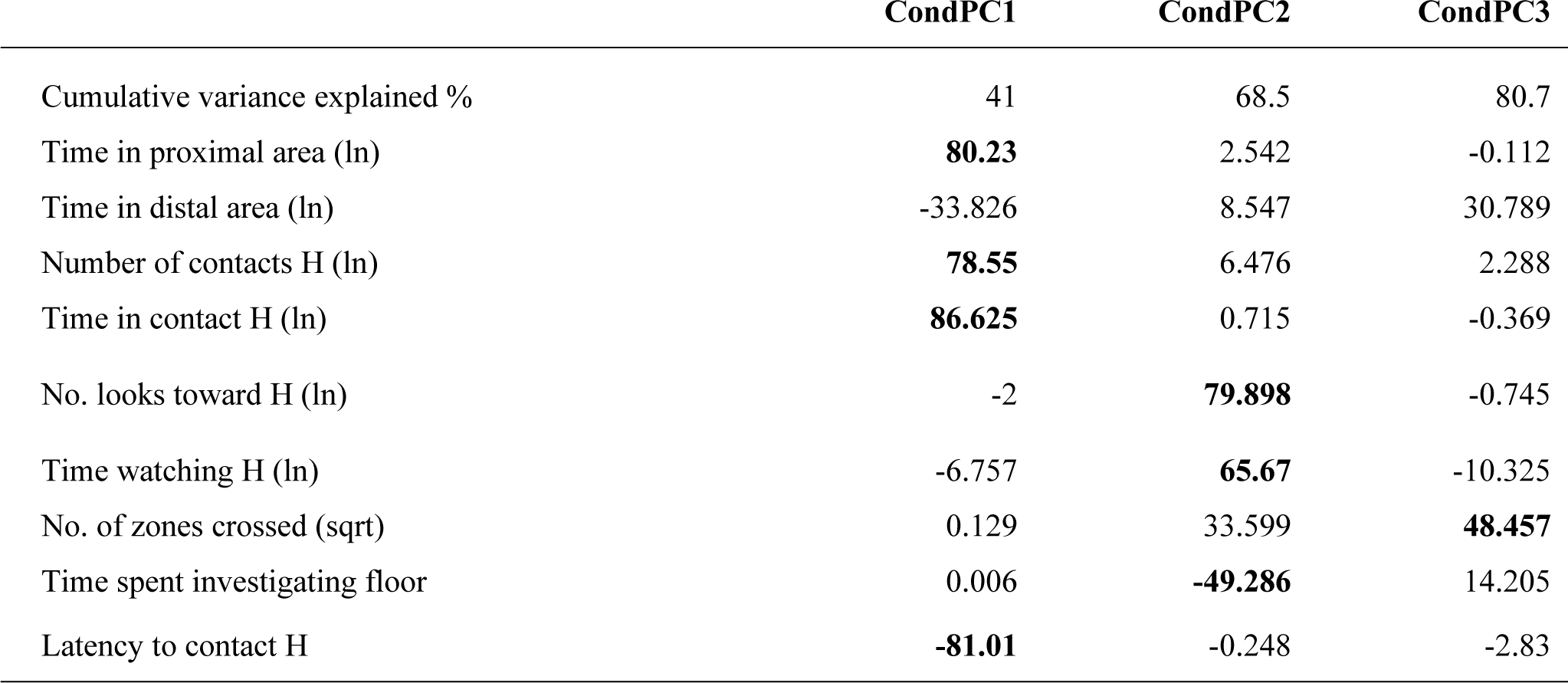
Percentage of explained variance and relative loadings of parameters on PCs, following the Principal Component Analysis computed on the behaviours scored during the sessions of additional positive contacts of the conditioning. The first three PCs, having an eigenvalue above 1 constituted three behavioural scores: CondPC1, CondPC2, CondPC3. Parameters that explain the most each PC are bolded (|loading|>0.4).

|  | CondPC1 | CondPC2 | CondPC3 |
| --- | --- | --- | --- |
| Cumulative variance explained % | 41 | 68.5 | 80.7 |
| Time in proximal area (ln) | **80.23** | 2.542 | -0.112 |
| Time in distal area (ln) | -33.826 | 8.547 | 30.789 |
| Number of contacts H (ln) | **78.55** | 6.476 | 2.288 |
| Time in contact H (ln) | **86.625** | 0.715 | -0.369 |
| No. looks toward H (ln) | -2 | **79.898** | -0.745 |
| Time watching H (ln) | -6.757 | **65.67** | -10.325 |
| No. of zones crossed (sqrt) | 0.129 | 33.599 | **48.457** |
| Time spent investigating floor | 0.006 | **-49.286** | 14.205 |
| Latency to contact H | **-81.01** | -0.248 | -2.83 |

**Table 4:**
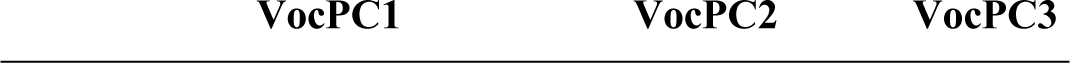
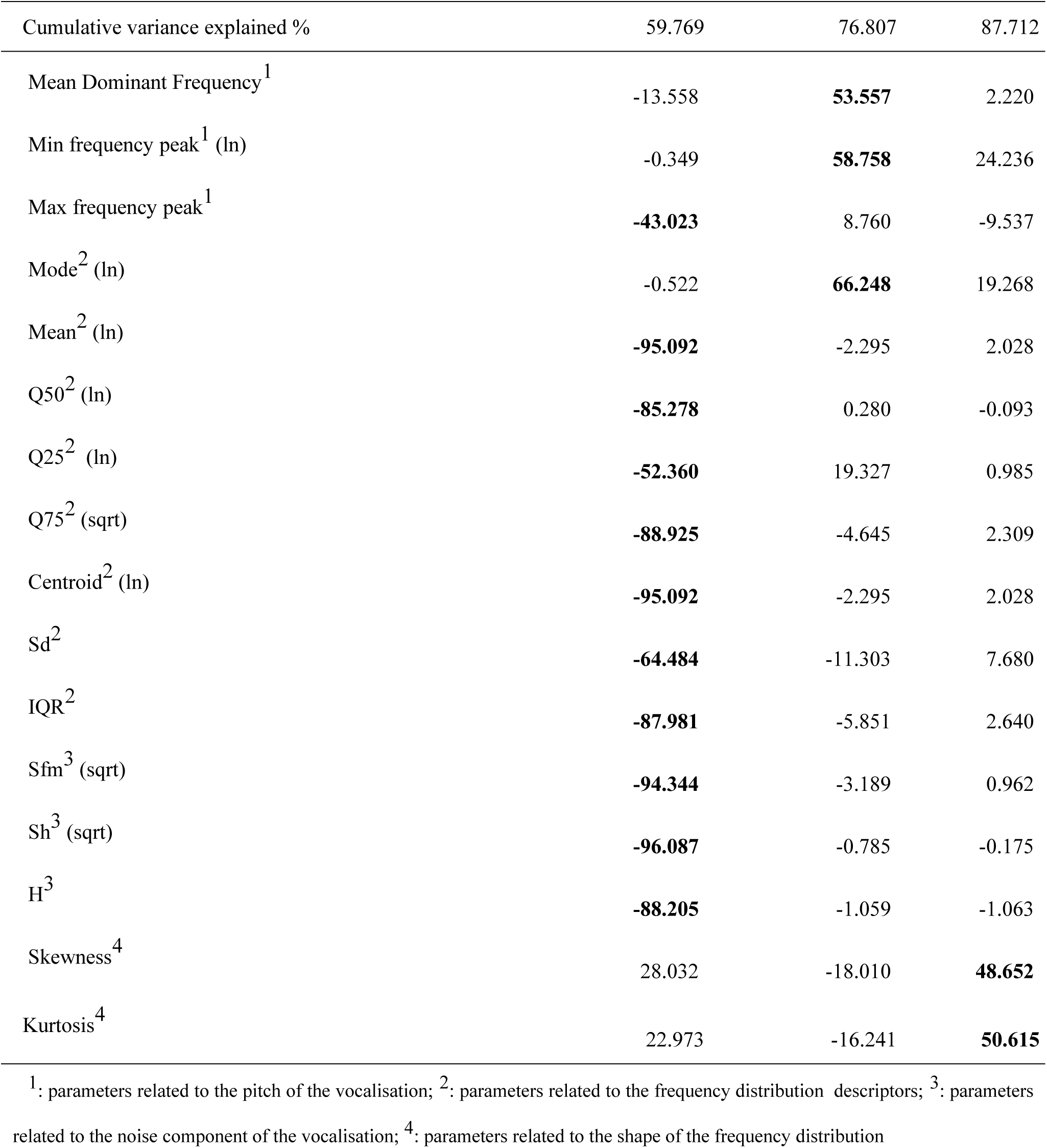
Percentage of explained variance and relative loadings of parameters on PCs following a Principal Component Analysis on spectral parameters of the grunts recorded in the entire dataset (including both types of tests, N=17 546 grunts). The transformations used to reach symmetrical distribution before the PCA are indicated in parentheses. The first three PCs, having an eigenvalue above 1 constituted three vocal response scores: VocPC1, VocPC2, VocPC3. Parameters that explain the most each PC are bolded (|loading|>0.4) .

##### Statistical models

All statistics were carried out on R (R Core Team, 2022). Linear mixed effect models [‘lmer’ function, ‘lme4’ R package (Bates et al., 2014)] were built when tested variables were linear (behavioural and vocal scores, grunt duration) and one binomial generalized mixed effect model was built for binary parameters (occurrence of missed contacts initiated by human during the conditioning). The following subsections describe how models were built for each type of tests. In all models described below, the identity of the replicate (‘1’ or ‘2’) was used as an interacting fixed factor, since the experiment was run in two identical replicates on two independent groups. The identity of the human (‘AH’ or ‘AV’) was used as interacting fixed factor in all models described below, since two experimenters were involved in the positive handling at weaning and in the session of additional positive contacts of the conditioning (but always the same human was attributed to a given piglet). The piglet was used as random factor to take into account the within-subject design. All explanatory variables used in the models and interactions between them were built in respect to the experimental design and to allow biological interpretations. As a consequence, not all interactions between all variables were made. They are fully explained in the subsequent sections.

##### Isolation/Reunion tests

The aim of this part was to test the effect of the positive handling at weaning treatment (H vs. H+ piglets) and additional human contacts during sessions of the conditioning on the piglet’s reaction to human presence. Since the same Isolation/Reunion test was repeated before and after the conditioning, we used the variable ‘Conditioning time’ as a two level interacting factor (‘before’ or ‘after’ conditioning, referred as “Time” in the models) to test the effect of the conditioning. Piglets spacial behaviour and proximity to the human was studied only during the reunion phase with the human that followed the isolation phase. Model_1 was computed:

Model_1 <– lmer (ReuPCs ∼ Treatment*Time + Treatment*Replicate + Treatment*HumanID + Time*Replicate + Time*HumanID + (1 | pigletID), data= data_Behaviour_Reunion).

Concerning the analysis of vocal behaviour, the isolation phase represents a negative social context for the piglets and may be used as a negative control when monitoring the effect of human presence on vocal expression of emotional states (Villain, Lanthony, et al., 2020). So, the two phases of the test were used to study the three way interaction between treatment (H vs.. H+), phase of the test (isolation vs.. reunion) and time of the conditioning (before vs.. after). The following model_2 was computed:

Model_2 <– lmer (VocPCs ∼ Treatment*Phase*Time + Treatment*HumanID + Time*HumanID

+ Treatment*Replicate + Time*Replicate + (1 | pigletID/Time/Phase), data= data_Vocal_Isolation + data_Vocal_Reunion).

To go further, only the reunion phase was kept and a proximity variable was added. Indeed, the piglet could vocalise either when close to human or away from them and this spatial proximity was demonstrated as an important factor of changes of vocal features (Villain et al. 2020b). Thus, a two level proximity factor was built: either ‘1’ when the piglet was in the proximal area (figure 1) or ‘0’ when it was elsewhere in the room. The following model_3 was computed:

Model_3 <-lmer (Vocal response score ∼ Treatment*Time*InProxArea +

###### Treatment*HumanID + InProxArea*HumanID + Treatment*Replicate + InProxArea*Replicate + Time*Replicate + Time*HumanID + (1 | pigletID/Time), data

= data_Vocal_Reunion).

##### Conditioning trials

The aim was to study the evolution of human-piglet relationship over the conditioning [the variable ‘Trial number’, used as a continuous variable, referred as “Trial” in the models]. The effect of treatment (positively handled at weaning H+ piglets or non handled H piglets) was tested as an interacting factor with Trial. Trial was also used as a random slope to take into account individual trajectories (Schielzeth and Forstmeier 2009). The following model_4 was built to test the behavioural response scores CondPCs (lmer) and the occurrence of missed contact initiated by the human during a session (presence/absence, binomial model, glmer):

Model_4 <– (g)lmer (CondPCs / Missed contact ∼ Trial*Treatment + Trial*HumanID + Trial*Replicate + Treatment*Replicate + Treatment*humanID + (1+ Trial | pigletID), (family=Binomial), data= data_Behaviour_Conditioning).

For the analysis of vocal response scores, similarly to the Isolation/Reunion test, the piglet could vocalise either when close to the human or away from them. We thus added the proximity factor in the analysis of vocal response variables. The following model_5 was built :

Model_5 <– lmer (VocPCs ∼ Trial*Treatment*InProxArea+ Trial*HumanID + Trial*Replicate + Treatment*Replicate + Treatment*HumanID + HumanID*InProxArea + Replicate*InProxArea + (1+ Trial | pigletID), data= data_Vocal_Conditioning).

##### Model validation and statistical tests

All linear models were validated by visual inspection of the symmetrical and normal distribution of the residuals. Anovas (‘car’ R package (Fox & Weisberg, 2011)) were computed on models to test for significant effects of explanatory variables. Following the Anova, when interactions were found significant, post hoc test were run on model interactions, correcting for multiple testing with Tukey contrasts (‘emmeans’ or ‘lstrends’ functions from ‘emmeans’ R package (Lenth, 2016), for categorical or continuous variables respectively). Considering the conditioning time (before or after conditioning), when involved in a significant three-way interaction, this factor was fixed to allow pairwise comparison within each time period as it was not considered relevant to assess the effect of time only. Results of the Anova, model estimates and pairwise post hoc comparisons are reported in the supplementary material (tables S1 and S2 for tests, table S3 for model estimates).

## Results

### Effect of positive handling at weaning and conditioning on piglets’ reaction to human presence (Isolation/Reunion tests)

#### Piglets that were not handled at weaning express a similar behavioural proximity to a human after a positive conditioning as the positively handled ones

The interaction between the treatment (positively handled piglets at weaning (H+) or not (H) and the conditioning time (before or after the conditioning) was significant for both ReuPC1 and ReuPC3 (𝜒^2^_1_ = 28.0, p < 0.001, and 𝜒^2^_1_ = 3.7, p = 0.05 respectively, figure 2) but not for ReuPC2 (𝜒^2^_1_ < 0.001, p = 0. 99, supplementary table S1). Post hoc tests on ReuPC1 showed that ReuPC1 was higher after the conditioning than before (H: after – before, t.ratio = 12.1, p <0.001, H+: after – before t.ratio = 11.0, p < 0.001) and that before the conditioning, piglets that were positively handled at weaning had significantly higher ReuPC1 than non handled piglets (Before, H – H+: t.ratio = -2.1, p < 0.001), but not after (After, H – H+: t.ratio = 0.02, p = 1.0). According to the loadings, this means that piglets that were positively handled at weaning had a lower latency to contact the human, approached them more often and spent more time close to and investigating the human (ReuPC1) than non handled piglets, before the conditioning. This score increased after the conditioning and no evidence of a difference between treatments after the conditioning was found (figure 2). Post hoc tests on ReuPC3 showed a significant effect of the conditioning time only in piglets that were positively handled at weaning (H+: after – before, t.ratio = 5.2, p < 0.001, H: after – before, t.ratio = 2.6, p = 0.06). No difference in ReuPC3 was found between treatments before the conditioning (Before: H – H+, t.ratio = -0.75, p = 0.87), whereas positively handled piglets had a higher -ReuPC3 after the conditioning than before (After : H – H+, t.ratio = -3.2, p = 0.009). According to the loadings, this means that after the conditioning, piglets that were positively handled at weaning expressed more investigation of the room after the conditioning than non handled piglets. No evidence of any effect on ReuPC2 was found (table S2).

**Figure 2:**
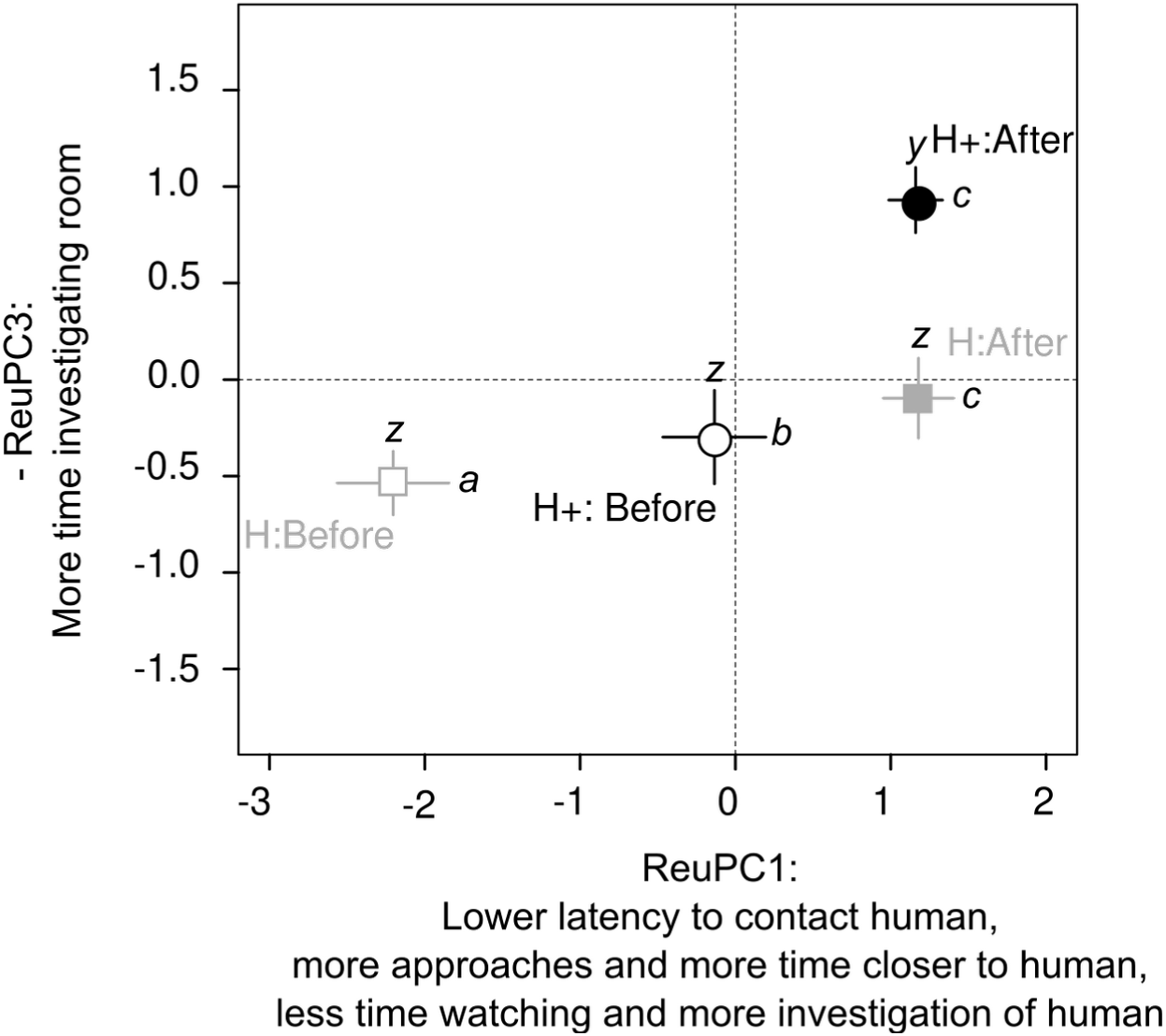
Effect of conditioning and treatment on spatial behaviour and proximity toward the human during the reunion of the Isolation/Reunion test. Mean ± SE per group is indicated, different letters indicates significantly different groups. Significant interaction between treatment (H : grey squares and H+ : black circles) and time (Before the conditioning: empty elements and After the conditioning: filled elements) on behavioural ReuPC1 (letters a to c) and ReuPC3 (letters z and y). Full statistical report is available as supplementary material (tables S1 S2 for statistical tests and S3 for model estimates)

#### Piglets positively handled at weaning produce shorter grunts even when no human is present

Using the isolation phase as a negative control we could compare the effect of the phase of the test (Isolation vs. Reunion with the human), taking into account the conditioning time (before or after the conditioning) and the treatment. No evidence of any effect of neither the three way interaction (𝜒^2^_1_ < 0.62, p > 0.43) nor two way interactions of interest was found (treatment: phase, conditioning time:phase, conditioning time: treatment interactions : 𝜒^2^_1_ <3.5, p > 0.06, table S2) in any of the scores.

Regardless of the treatment, single effects of the phase of the test were significant for grunt duration and all AcPCs (𝜒^2^_1_ > 6.6, p < 0.01, table S1). During the reunion phase with the human, grunts were shorter (estimates of log(duration)[95% CI] : -1.32[-1.37;-1.26] vs. -1.06[-1.12;-1.00]), had a higher frequency range, higher bandwidth and a higher noise component (-VocPC1: 0.78[0.48;1.08] vs. 0.34[0.03;0.66]), were higher pitched (VocPC2: -0.18[-0.36;0.01] vs. -0.46[-0.65;-0.28]) and their spectrum had a higher skewness and kurtosis (VocPC3: -0.25[-0.37;-0.14] vs. - 0.11[-0.23;0.01]), compared to the isolation phase.

Regardless of the phase of the test, single effects of treatment were found for grunt duration and - VocPC3 (𝜒^2^_1_ = 5.5, p = 0.02 and 𝜒^2^_1_ = 4.9, p = 0.03 respectively, table S2). Grunts produced by positively handled at weaning piglets were shorter (estimates of log(duration)[95% CI]: -1.25[-1.32;-1.19] vs. -1.12[-1.2;-1.1], table S3), and differed in -VocPC3 scores, describing the shape of the frequency spectrum (estimates of -VocPC3[95% CI]: -0.29[-0.43;-0.14] vs. -0.07[-0.22;0.08], table S3), than grunts produced by non handled piglets.

#### Positive handling and conditioning affect vocal reactivity to human proximity

During the 5 min reunion, the piglet was scored either as close to the human or away from them. The three way interaction of the conditioning time, the treatment and the location was significant for grunt duration, -VocPC1 and VocPC3 (𝜒^2^_1_ > 4.9, p < 0.03). Post hoc tests revealed that grunts produced closer to the human were shorter than the ones produced further away, but only in piglets that were not positively handled at weaning, effect being stronger before the conditioning than after it (H piglets: away – close, z.ratio = 6.3, p < 0.001 before and z.ratio = 4.1 p < 0.001 after the conditioning; H+ piglets: away – close z.ratio < 1.98 p > 0.19, figure 3A). -VocPC1 was higher, i.e. grunts had a higher frequency range, bandwidth and were noisier when produced closer to the human than further away, but only in non handled piglets and before the conditioning (H piglets: away – close, z.ratio = -3.34, p = 0.005 before and z.ratio = -1.23 p = 0.61 after the conditioning; H+ piglets: away – close, z.ratio < 0.36 p > 0.21, figure 3B). For VocPC2, the three way interaction did not reach significance (𝜒^2^_1_ = 3.3, p = 0.07), so only subsequent two way interactions were considered (post hoc tests on the three way interaction can be found in supplementary, tables S1 to S3). For VocPC2, significant two way interactions were found between the conditioning time and the location (𝜒^2^_1_ = 10.3, p = 0.001) on the one hand, and between the location and the treatment (𝜒^2^_1_ = 4.2, p = 0.04) on the other hand. Post hoc tests revealed that grunts produced closer to the human had a higher VocPC2, meaning they had a higher pitch, effect being stronger before the conditioning than after (before: away – close, z.ratio = -6.12, p < 0.001; after: away – close, z.ratio = -2.88, p = 0.004, figure 3C). The increase in VocPC2 with the location was greater for non handled piglets than positively handled piglets (H piglets: away – close, z.ratio = -5.54, p < 0.001; H+ piglets: away – close, z.ratio = -3.82, p = 0.001, figure 3D). The last two-way interaction of interest between the conditioning time and the treatment did not reach significant level (𝜒^2^_1_ = 0.80, p = 0.37). For VocPC3, post hoc tests did not reach significant levels (|z.ratio| < 2.3 p > 0.09 for any comparison) .

**Figure 3:**
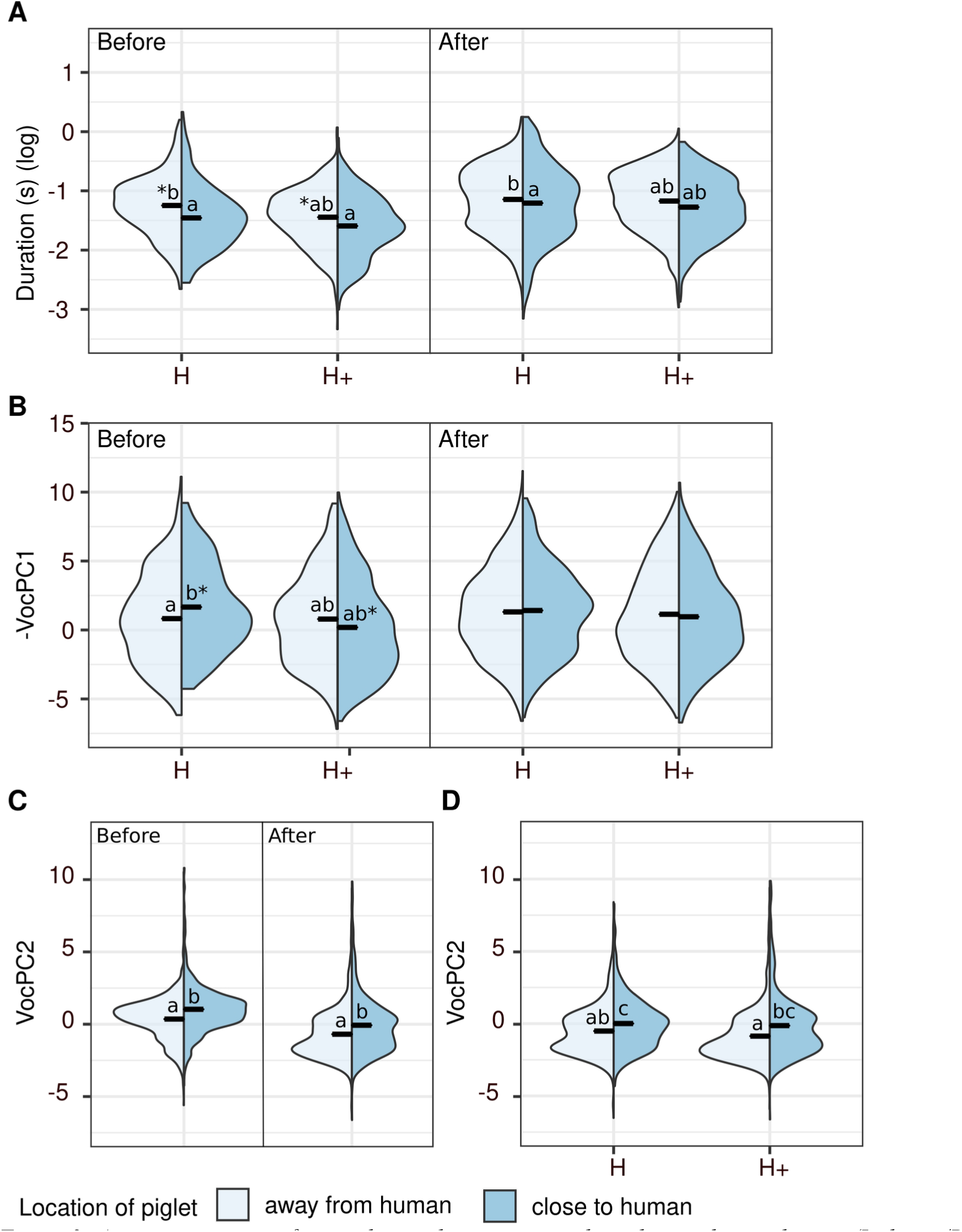
Acoustic structure of grunt during the reunions with a silent and static human (Isolation/Reunion test). Effect of conditioning (before or after), treatment (H or H+) and location of the piglet relatively to the human (close: dark blue or away from them: light blue). Violin plots representing the median and the density of data distribution in the considered groups. (A, B) Results of post hoc tests following significant three way interaction between treatment, conditioning time and location on grunt duration (A) and on the first vocal score -VocPC1 (B). (C,D) Results of post hoc tests following significant two way interactions between conditioning time and location (C) and between treatment and location (D) on the second vocal score VocPC2. Values with no common letters differ significantly. When no letters are present, no significant difference between groups was found. Stars (*) between two groups represent a statistical trend (p< 0.10). Full statistical report is available as supplementary material (tables S1 S2 for statistical test and S3 for model estimates).

### Emergence of positive perception of human (effect of additional positive contacts sessions over the conditioning)

#### The conditioning increases behavioural proximity to the human in all piglets

No evidence of any effect of the interaction between the treatment [positively handled piglets before the conditioning (H+) or not (H)] and the trial number was found for all behavioural scores (CondPC1, CondPC2 and CondPC3, table 3). Independently from the treatment, the higher the trial number the higher CondPC1 (𝜒^2^_1_ = 59.3, p < 0.001, slope estimate [95% confidence interval]: 0.20 [0.15 : 0.25]) and the lower CondPC2 was (𝜒^2^_1_ = 48.6, p < 0.001, slope estimate: -0.17 [-0.22 : - 0.12]). According to the loadings, over the conditioning, piglets decreased the latency to contact the human, made more contacts, spent more time in the proximal area and in contact with the human (condPC1), decreased the number of looks to the human, spent less watching the human and more time investigating the room (CondPC2) (figure 4A). Independently from the trial number, positively handled piglets had a lower CondPC2 and a higher CondPC3 than the non handled ones (𝜒^2^_1_ = 12.8, p < 0.001 and 𝜒^2^_1_ = 7.0, p = 0.008 respectively), meaning that piglets that were positively handled at weaning expressed a fewer number of looks to the human, spent less time watching them and more time investigating the room (CondPC2) and crossed more virtual zone during the test (CondPC3) (figure 4B). The probability of having at least one missed contact by the human during a session was lower for positively handled piglets than non handled ones (𝜒^2^_1_ = 9.57, p = 0.002, figure 4C), with no interaction with the trial number (𝜒^2^_1_ = 0.22, p = 0.064).

**Figure 4:**
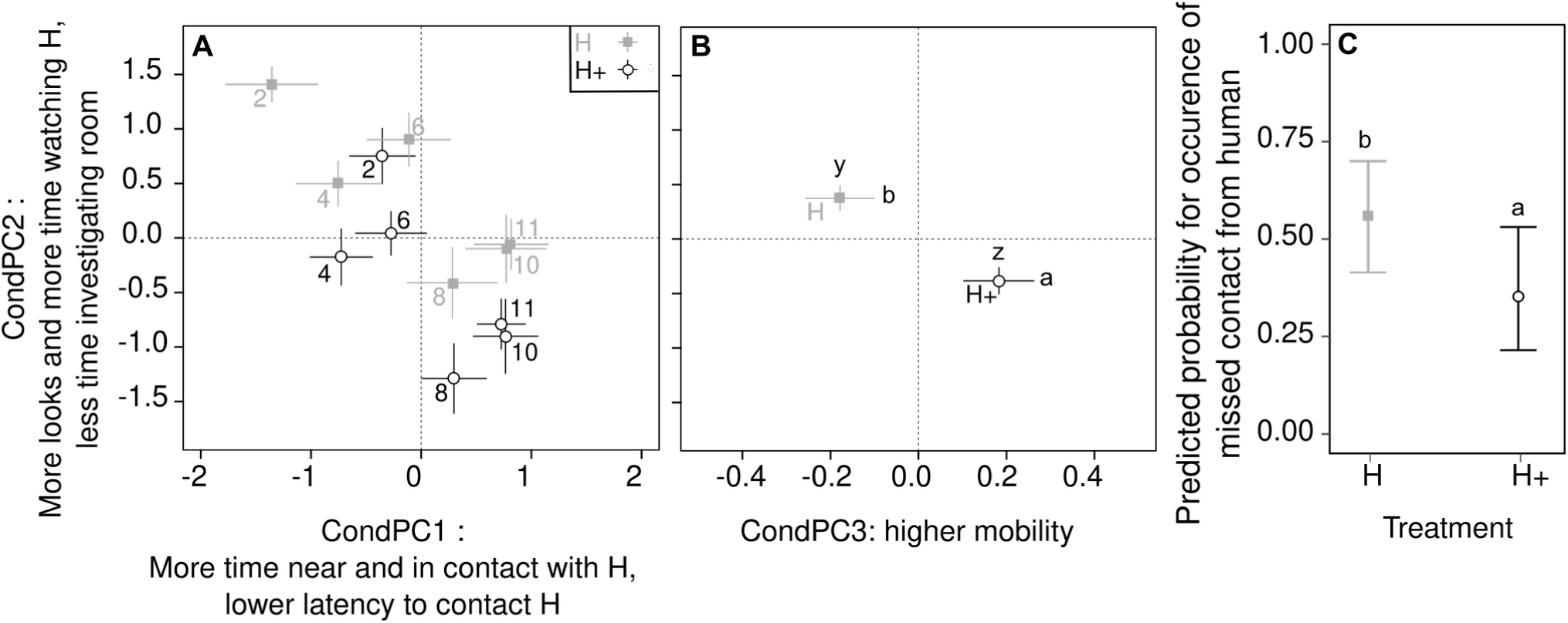
Behavioural variation of responses of piglets according to the sessions of additional positive contacts of the conditioning (A), and to the treatments (B, C). (A,B) Mean ± SE per group. Numbers in (A) refers to the trial number of the conditioning. Higher CondPC1 and lower CondPC2 over time (single effect of trial number, A). Higher CondPC2 in H piglets than H+ piglets regardless of time (single effect of treatment, A). Higher CondPC3 and lower CondPC2 in H+ piglets than H piglets (single effect of treatment, B). (C) Mean estimates ± 95% confidence interval from generalized mixed effect model. Lower probability of occurrence of missed contact by the human in H+ than H piglets (single effect of treatment). Full statistical report is available as supplementary material (tables S1 et S2 for statistical tests, table S3 for model estimates).

#### Additional positive contacts trigger shorter and higher pitch grunts in all piglets

During the sessions of additional positive contacts of the conditioning, the three-way interaction between the trial number, the treatment and the location was not significant for any of the vocal scores (𝜒^2^_1_ < 0.18, p > 0.67), allowing the analysis of the two way interactions of interest. The interaction between treatment and the trial number was not significant for all vocal scores (𝜒^2^_1_ < 2.5 p > 0.11). Grunt duration decreased over time and independently from the treatment (trial number:replicate interaction, 𝜒^2^_1_ <5.3 p = 0.02, slope estimate -0.03[-0.04;-0.01] for the lower slope, table S1 and S3). However, independently from the trial number, grunt duration was lower when piglets were located close to the human and this effect was stronger in non handled piglets than positively handled piglets (treatment:location interaction: 𝜒^2^_1_ = 15.8 p < 0.001, away vs.. close, H piglets: z.ratio = 10.2 p < 0.001, H+ piglets: z.ratio = 6.86 p < 0.001, figure 5A). -VocPC1 and VocPC2 decreased over time but remained higher when piglets were located close to the human (trial number: location interaction, 𝜒^2^_1_ = 3.97 p = 0.05 and 𝜒^2^_1_ = 6.1 p = 0.01 respectively for -VocPC1 and VocPC2). According to the loadings, this means that the frequency range, bandwidth and noisiness of grunts (-VocPC1) as well as the pitch (VocPC2) decreased over the conditioning when piglets were located away from the human but remained high when piglets were close (slope comparison away – close, -VocPC1 : z.ratio = -1.80 p = 0.07, VocPC2 : z.ratio = -2.34 p = 0.02, figure 5C). Additionally, VocPC2 was higher when piglets were close to the human in non handled piglets (treatment:location interaction, 𝜒^2^_1_ = 7.6 p = 0.005, pairwise comparisons away vs. close, in H: z.ratio = -4.9 p z 0.001 and in H+: z.ratio = -2.0 p = 0.21), meaning that non handled piglets produced higher pitched grunts when closer to the human (figure 5B).

**Figure 5:**
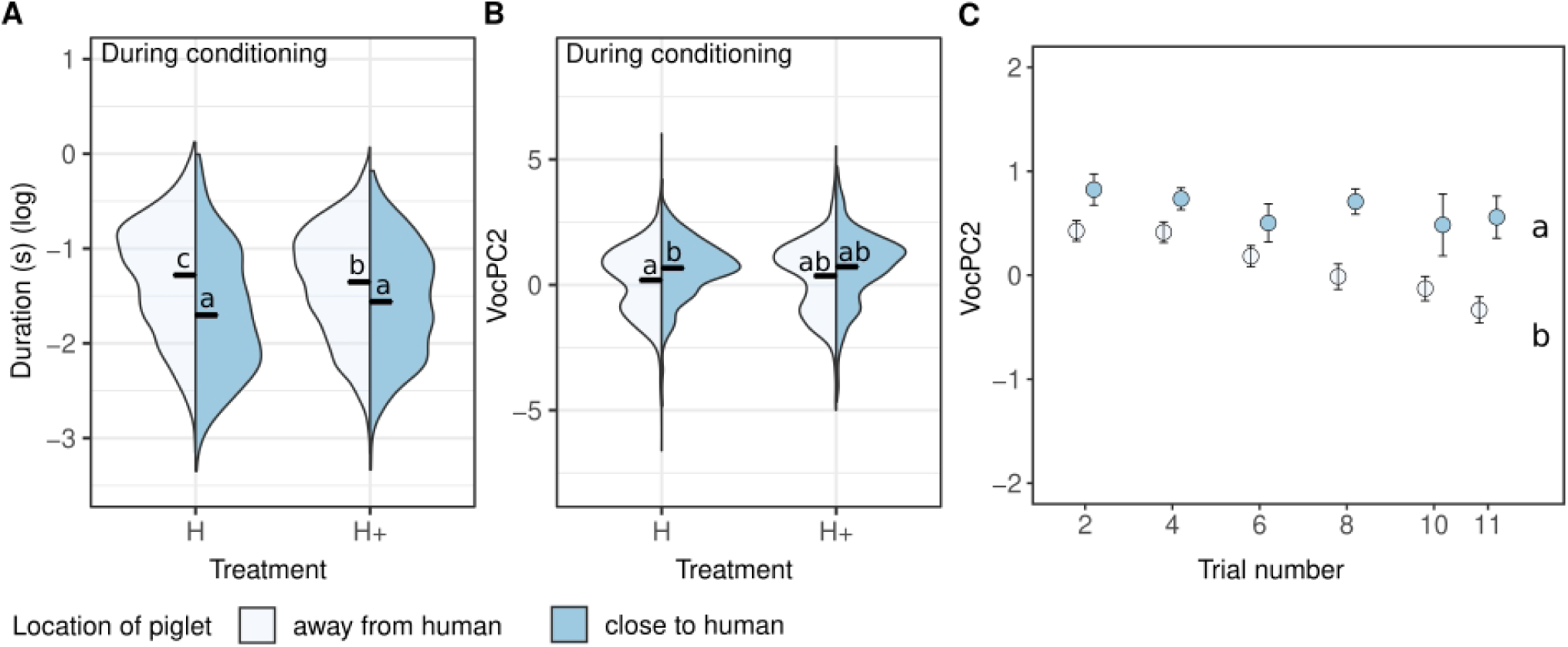
Vocal scores over the conditioning, during the 2min sessions of additional positive contacts. (A, B) Violin plots representing the median and the density of data distribution in the group. Interacting effect of location (in proximal area of the human ‘(close’: dark blue) or elsewhere in the room (‘away’ from the human: light blue) and treatment (H vs. H+ piglets) on grunt duration (A) and VocPC2 (B). (C) Mean ± SE per group, interacting effect of trial number and location of piglets on VocPC2. Values with no common letters differ significantly (difference between groups: A, B or slopes: C). Full statistical report is available as supplementary material (tables S1-S3).

### Impact of human identity on piglets behaviour and grunt structure

Since half of the piglets had been assigned to one human experimenter and the other half to another one, the identity of the human was included in the model. This allowed to test interactions between the identity of the human and the treatment of positive handling at weaning on the one hand and the conditioning time on the other hand.

During the reunions of the Isolation/Reunion test, the interaction between treatment and human identity was significant for the first behavioural proximity score (ReuPC1, 𝜒^2^_1_ = 6.01, p = 0.01) but not the others (ReuPC2 and ReuPC3 (𝜒^2^_1_ < 1.98, p > 0.16, table S1). The effect of treatment on ReuPC1 was higher when piglets were handled by the human ‘AH’ (H vs. H+, AH: t.ratio = -4.77, p < 0.001, figure 6). When the human ‘AV’ handled the piglets, for which ReuPC1 scores exhibited intermediate values, treatment was not significant (AV, H vs. H+: t.ratio = -1.33, p = 0.56). These interacting effects of the human identity and treatment on behaviour were not found when considering the reunions of the conditioning (𝜒^2^_1_ < 1.32, p > 0.25 for all CondPCs, table S1).

**Figure 6:**
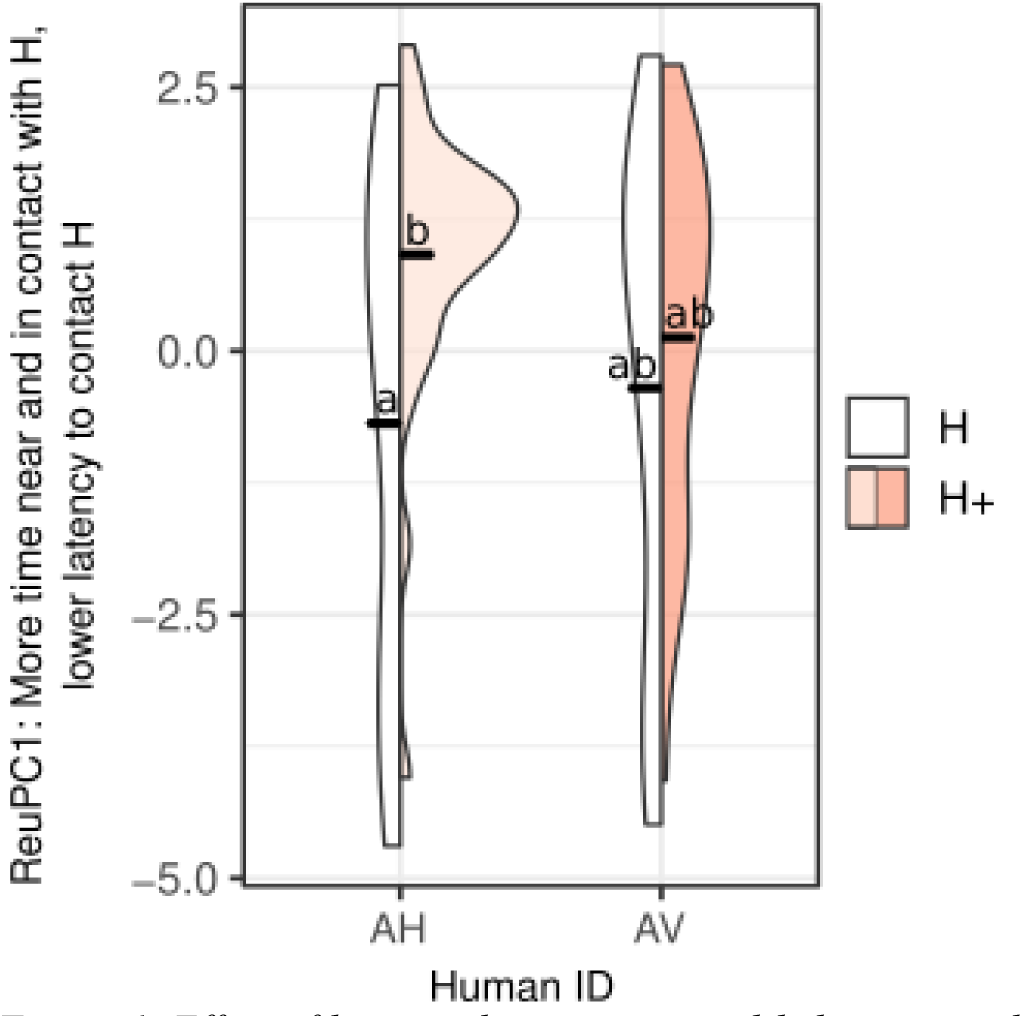
Effect of human identity on spatial behaviour and proximity during the reunion of the Isolation/Reunion test. Violin plots representing the median and the density of data distribution in the group. Values with no common letters differ significantly. Full statistical report is available as supplementary material (tables S1 and S2 for statistical tests, table S3 for model estimates).

Interactions between the human identity and conditioning time were not significant, neither considering the reunions of the Isolation/Reunion test (ReuPCs, 𝜒^2^_1_ < 0.642, p > 0.42, tables S1), neither the trial number during the session of additional positive contacts of the conditioning (CondPCs, 𝜒^2^_1_ < 0.11 p > 0.74, table S1).

Considering the vocal scores, no effect of human identity was found on VocPC1 during the Isolation/Reunion tests but -VocPC1 was higher when the human ‘AH’ was in the room during the reunion periods of the conditioning (table 5), meaning the frequency range and the bandwidth of the grunt were higher when the human ‘AH’ interacted with the piglet compared to the human ‘AV’. VocPC2 was higher when the human ‘AH’ was in the room during the Isolation/Reunion tests (table 5), meaning that the pitch of grunts was higher and this effect was also found during the sessions of additional positive contacts of the conditioning in interaction with the location of the piglet (𝜒^2^_1_ = 11.9, p = 0.001): VocPC2 increased when piglets were located close to the human but this increase was significant only for the human ‘AV’ and not for ‘AH’ (table 5). VocPC3 was not different between humans during the reunions of the Isolation/Reunion tests but, over the conditioning, VocPC3 changed differently when piglets were handled by the human ‘AH’ or ‘AV’, as showed by the significant interaction between trial number and human identity (𝜒^2^_1_ = 8.0, p = 0.005): the skewness and kurtosis of grunts decreased over the conditioning when ‘AH’ was interacting with the piglets, but not ‘AV’ (see slope estimates, table 5). No evidence of any effect of human identity was found on grunt duration neither during the Isolation/Reunion tests nor during the sessions of additional positive contacts of the conditioning (table S1).

**Table 5:** Significant effects of human identity on vocal response score (VocCP1 and VocPC2) during the reunion of the Isolation/Reunion test and during the sessions of additional positive contacts of the conditioning. Only significant effect are presented here but a full statistical report is available as supplementary material (tables S1 and S2 for statistical tests, table S3 for model estimates). When single effects were interpretable, the Chi-squared statistic are reported. When significant interactions were significant, post hoc pairwise comparisons were performed with Tukey corrected and are thus reported. The estimates correspond either to the group estimate and comparisons of groups (categorical fixed effect) or slope estimates and comparison of slopes (continuous fixed effect, ‘Trial number’).

| Vocal response score | Fixed effect | Levels | Estimate | Lower.95%CI | Upper.95%CI | Statistic | P-value |
| --- | --- | --- | --- | --- | --- | --- | --- |
| Reunion of the Isolation/Reunion test |  |  |  |  |  |  |  |
| VocPC2 | humanID | AH | 0.154 | -0.119 | 0.427 | $\chi^2_1 = 4.94$ | P = 0.03 |
|  |  | AV | -0.292 | -0.571 | -0.012 |  |  |
| Sessions of additional positive contacts of the conditioning |  |  |  |  |  |  |  |
| -VocPC1 | humanID | AH | 0.091 | 0.037 | 0.144 | $\chi^2_1 = 4.69$ | P = 0.03 |
|  |  | AV | 0.076 | 0.021 | 0.132 |  |  |
| VocPC2 | humanID*I<br>nProxArea | AH – away | 0.317 | 0.110 | 0.524 | Z-ratio = -1.23 | P = 0.60 |
|  |  | AH – close | 0.402 | 0.161 | 0.643 |  |  |
|  |  | AV – away | 0.027 | -0.182 | 0.236 | Z-ratio = -5.77 | P < 0.001 |
|  |  | AV – close | 0.462 | 0.212 | 0.712 |  |  |
| VocPC3 | humanID *<br>Trial number | AH – Trial<br>number | -0.048 | -0.070 | -0.026 | Z-ratio = -2.82 | P = 0.005 |
|  |  | AV – Trial<br>number | -0.007 | -0.031 | 0.016 |  |  |

## Discussion

In this study, familiarity to a human and human-animal interactions were experimentally modified in weaned piglets to study the establishment of a positive HAR and test whether grunt structure could reflect a positive HAR. A positive conditioning paradigm, using additional positive contacts from a human as a reward, allowed to compare the behavioural changes over time in piglets previously positively handled at weaning or not. Two types of sessions were studied: a standard isolation/reunion tests with the human, carried out before and after conditioning, during which the human remained silent and did not interact with the piglet, and sessions of the conditioning, during which the human interacted with the piglets, providing additional positive contacts, as long as the piglets stayed close to the seated human. Behavioural data were collected to describe the positive HAR. Grunts produced during the tests and sessions were collected and their spectro-temporal structure confronted to the behavioural data, with the hypothesis that vocalisation structure may reflect the quality of HAR, though vocal markers of positive emotions. Firstly, the discussion will focus on the behavioural validation of the establishment of a positive HAR. Secondly, behavioural and vocal expression will be confronted to discuss grunt spectro-temporal structure as indicator of the quality of HAR. Last, we will discuss perspectives regarding the effect of human identity on the establishment of a positive HAR.

### Behavioural evidence of a rapid establishment of interest and proximity toward a human providing additional positive contacts

The standard reunion test with the human before the conditioning showed first that the treatment of positive handling at weaning succeeded in creating two different levels of human-piglet relationship (H and H+), as positively handled piglets expressed a higher attraction toward the human than non handled piglets (ReuPC1), parameters considered as indicators of a positive HAR (Rault et al., 2020). Second, this test showed that the conditioning increased the behavioural proximity toward the human of both positively handled and non handled piglets so that non handled piglets expressed a similar attraction toward the human as positively handled piglets. These results are in line with the behavioural results of the sessions of additional positive contacts. The analysis of piglets’ behaviour every second sessions of the conditioning showed that, although positively handled and non handled piglets started with different degree of proximity toward the human (trials 2 and 4, CondPC1), then, over time and for both treatments (H and H+), piglets expressed a higher attraction toward the human (CondPC1) and avoided less the human when the latter attempted to interact with them. So it seems that the conditioning process allowed non handled piglets to compensate the lack of positive handling before the conditioning and develop a similar proximity toward the human. Two minute daily sessions of additional positive contacts changed positively the perception of the human for the piglets, and thus their willingness to interact with them. Since no evidence of any interaction between time and treatment was found, no conclusion on differential developmental trajectories between treatments can be drawn, but a parallel development of the human-piglet relationship in both groups, when considering the proximity.

Beside behavioural proximity, piglets that were positively handled at weaning expressed more exploratory behaviours than non handled piglets after the conditioning (ReuPC3). This was also observed during the sessions of additional positive contacts of the conditioning: positive handled piglets started with a higher score associated with investigation than non handled piglets (CondPC2) and it held over the conditioning. Piglets that were positively handled at weaning also expressed a higher mobility than non handled piglets (CondPC3). These observations may be interpreted as an expression of natural foraging and disinterest from human contact, which may be a sign of positive welfare (Weerd & Day, 2009). In addition, this could also be interpreted in terms of attachment to the human. Indeed, attachment to a human may facilitate exploration of novel environments or objects, as shown in dogs (Palmer & Custance, 2008). A period of positive handling at weaning may provide an environment secure enough for the piglets to explore their environment in the presence of the human. Attachment has also been hypothesised in the lambs-human relationship (Tallet et al., 2009).

Overall, the behavioural monitoring showed that 2 min sessions of positive additional contacts per day are sufficient to increase proximity to a human to similar levels as when piglets were previously familiarised for 2 weeks, even when piglets experienced social isolation. But it did not allow the non handled piglets to express natural exploratory behaviours as the positively handled piglets. We hypothesize a sequential establishment of a positive HAR over time: firstly with a decrease of attentive state and an increase in proximity and accepted contacts, and secondly with a disinterest of human contacts and the expression of natural foraging behaviour. The latter may require a higher exposure time.

In the next paragraph we discuss to what extent changes in grunt spectro-temporal structure may reflect behavioural changes linked to the positive HAR over time.

### Links between vocal expression and positive HAR

#### A positive HAR is reflected by shorter grunts in presence and absence of a human

The social isolation phase of the Isolation /Reunion test, before any human entered the room, was associated with longer, lower pitched grunts with a downshifted frequency spectrum, whereas the reunion with a static human changed grunts structure to shorter, higher pitched with an upshifted frequency spectrum and this was observed in both handled and non handled piglets (H or H+) as well as before and after the conditioning. In terms of emotional indicators, similar changes in acoustic features of grunts were found in studies focusing on vocal markers of valence in pigs (Briefer et al., 2019, 2022; Friel et al., 2019; Villain, Hazard, et al., 2020), meaning that the reunion with a human, after a period of social isolation would be perceived as positive. However, this modulation of grunt structure was observed regardless of piglet experience with the human. It is possible that the reunion with an either neutral or familiar human, releasing piglets from total isolation could be perceived as positive by the piglets, as suggested in previous studies (Villain, Lanthony, et al., 2020).

In addition, and surprisingly, positively handled piglets produced shorter grunts than non handled piglets regardless of human presence. This was previously shown in another context (anticipation of (pseudo)social events independently from the type of partner) in the same groups of piglets (Villain, Hazard, et al., 2020). This may show that the period of positive handling at weaning modulated vocal expression in the long term, as this result was found both before and after the conditioning. On the one hand, a positive HAR establishes through successive positive experiences (Rault et al. 2020) and, on the other hand, HAR may have long term effects on behavioural expressions, as suggested by Brajon et al. (2015) using cognitive bias tests. We can thus hypothesize this may also be reflected in the way piglets vocalise, in general. In that case, we may have evidence of expression of another category of affect, moods, and not only emotional expression. Indeed, as suggested by Schnall (2010), although emotions are short-term affects triggered by an external stimulus, moods, on the other hand, may be experienced on a longer term and may not be attributable to a specific stimulus. Although emotions and moods do not rely on the same time scale, they may interact with one another, and more studies are needed to understand their effects on vocal expression.

#### A positive HAR affects vocal reactivity toward a static human

In a previous study, we showed that pigs vocalizing close to a human that previously had provided repetitive additional positive contacts produced shorter and higher pitch grunts, compared to when vocalizing away from the human (Villain, Lanthony, et al., 2020). Using the same type of test with positively handled at weaning and non handled piglets, before or after conditioning sessions with positive interactions, we can test the effect of positive handling on this modulation of grunt structure. Similarly to the previous study, during the standard reunion test (no contact from the human), piglets produced shorter and higher pitched grunts with an upshifted frequency spectrum when close to the human. It has to be noted that this effect was 1) stronger in previously non handled piglets than positively handled at weaning piglets and 2) stronger before the conditioning than after. In other words, the more familiar with the human associated with positive handling, the less reactive to human proximity.

These results may be interpreted according to the behavioural results we described earlier (fig. 2). We described that the proximity to the human was first increasing at the beginning of positive handling experiences (see H piglets, before vs. after conditioning) before reaching a maximum (see H vs. H+ piglets after conditioning) and that the most familiar piglets showed more exploratory behaviours (H+ after conditioning). The acoustic results during the standard reunion mirror the behavioural results from the same test. The least familiar piglets would vocally express the exploration of a neutral and static human and, as the familiarity with the human increases, the human may become part of their environment, explaining the lack of vocal reactivity when close to the static human.

In addition, we may also be facing ceiling effects in terms of vocal flexibility, which could also partly explain these results. We showed that positively handled piglets generally produce shorter grunts than non handled piglets, and that the shape of the frequency spectrum of these grunts was different. So the structure of their calls, in general is different. According to the source-filter theory of vocal production, vocal flexibility is constrained by the dimensions and functioning of the vocal apparatus (lung capacity, characteristics of the vocal folds, length and shape of the vocal tract, see (Taylor & Reby, 2010) and (Titze & Martin, 1998)). It is possible that the positive HAR developed by the positively handled piglets may have change their grunts structure to an extent that vocal flexibility is no longer quantifiable in the experimental design of this study.

#### Providing rewarding additional positive contacts triggers short and high pitched grunts

Contrary to the standard reunions with a static human, the human actively interacted with the piglets during the sessions of the conditioning, providing contacts and producing speech as long as the piglets remained close to the human. During these sessions and contrary to the standard reunions, grunts produced close to the human were shorter and higher pitched, regardless of the trial number of the conditioning and treatment. Although these effects were stronger in non handled piglets than positively handled piglets, they remained over time. We describe here two types of vocal reaction to human proximity, depending on the human behaviour. On the one hand, time decreased vocal reactivity to human proximity during a standard reunion with a static human. On the other hand, no evidence of a decrease in vocal reactivity to human proximity was found during sessions of additional positive contacts. This would mean that positive interactions with piglets consistently triggers the production of shorter and higher pitch grunts. These changes may be explained by the expression of a higher arousal state experienced by the piglets while being positively handled. Indeed, in the context of these sessions, the piglet could choose to approach and stay close to the human, which will provide positive contacts systematically. So the piglet may anticipate to receive positive contact and systematically being rewarded. When close to the human, observed changes in frequency distribution of grunts (increased pitch and upshifted frequency spectrum) are known to be markers of arousal (in the negative state in multiple mamalian species (Briefer, 2012, 2020) and pigs (Linhart et al., 2015)). In addition, these spectral changes were also associated with shorter grunts. Although the duration of grunts is associated with the valence of a situation, the duration may also be an indicator of positive arousal. This hypothesis has to be taken precociously since no additional control of arousal could be done in the present study.

This working hypothesis may explain the decrease in vocal reactivity to human proximity observed during the standard reunion test as the HAR becomes more positive. Indeed, before the sessions of the conditioning, positively handled piglets were habituated to a human interacting positively when present whereas non handled piglets were not, hence, during the first standard reunion test, when the human is present but do not interact with the piglet, positively handled and non handled piglets may have diverging expectations regarding the presence of the static and silent human. As positively handled piglets received positive contacts every time they were in the presence of the human, they may have expected positive contacts when approaching and experienced an absence of reward during the test. This has already been hypothesised in piglets deprived from human voice during interactions after a period of habituation to it (Bensoussan et al. 2020). On the contrary, piglets that were not positively handled at weaning never experienced additional positive contacts and being close to a human, having the possibility to investigate them may be some kind of reward after the period of total isolation. After the conditioning, piglets from both treatments were conditioned to receive additional positive contacts and both groups had experienced a first standard reunion test, so they may both experience an absence of reward during the test, which may explain a lower reaction to human proximity, and thus fewer changes on grunt spectro-temporal features.

Last, we can raise the question whether changes in grunt structure in reaction to rewarding positive contacts may also be associated with a specific human-pig communication. In other domestic species, owner directed vocalisations has been shown (in cats, reviewed in (Turner, 2017); in dogs (Gaunet et al., 2022)). In addition, studies have found similar socio-communicative behaviours toward a human in socialized pigs and dogs (Gerencsér et al., 2019). Hence, we may profit from testing the existence of human directed vocalisations in pigs, as consequences of their socio communicative abilities.

### Effect of human identity on piglets’ perception: perspectives on HAR

We found that the identity of the human had effects on behavioural and vocal response scores. Piglets that were handled by the human ‘AH’ had higher values of behavioural proximity (ReuPC1) than piglets handled by the human ‘AV’ during reunion test after a period of isolation. This effect was not found during conditioning sessions. The effect of the human did not interact with the conditioning time, leading to the conclusion that the difference between the two experimenters may have established during the period of positive handling at weaning, prior to the conditioning. Additionally, when the human ‘AH’ was in the room, piglets produced grunts with a more upshifted frequency spectrum and a higher pitch than when the human ‘AV’ was in the room. If upshifted grunts may be a indicator of positive higher arousal, then we may conclude that ‘AH’ was more likely to trigger higher positive states than ‘AV’. Interestingly, the human identity and the spatial proximity had different effects on piglets grunts during sessions of additional positive contacts but not when the human was static during the standard reunion test. Hence, it is possible that the way one human interacts (behavioural and vocally) with a piglet may be more or less effective at triggering positive emotions and thus modifications of grunt structure. Several evidence exists in the literature that pigs discriminate humans visual and auditory cues (Bensoussan et al., 2019; Brajon et al., 2015c). Pigs may also show behavioural changes hearing human voice (Bensoussan et al., 2020). We may question the efficiency of different human features to generate a positive HAR. In our study, both humans that interacted with the piglets wear exactly the same clothes and standardized their tactile interactions toward the piglets before starting the study, and agreed on the rhythm and types of sounds (words, intonation) to use, to minimise generating variability although no systematic controls of the human behaviour or spectral feature of voices were performed here. It thus remains unclear whether experimenters interacted differently or if they were initially perceived differently by piglets. Our results show that the identity of the human may modulate piglet proximity and vocal behaviour but the design of this experiment does not allow to find the causes of these observations (behaviour, voice characteristics, or even odour profile). Thus, more studies of human features that are most likely to generate a positive HAR are needed and may be of interest regarding animal welfare. In addition, studying human-piglet relationship in a more systematic way, as in other domestic species, for example the play behaviour in dogs (Horowitz & Hecht, 2016) or the pet directed speech (Jeannin et al., 2017; Lansade et al., 2021), may shed light on the evolution and converging strategies of interspecific relationships. However, the influence of human identity did not modify the general outcomes of our study, but only decreased some effects, suggesting that this variability does not modify the main results, but should be considered in future studies.

To conclude, we showed that degrees of familiarity toward a human could be reflected in the way piglets vocalise in their presence, and out of it. We also showed that the spatial proximity toward a human providing additional care could change the acoustic structure of piglet grunts. These changes are likely to be linked to positive and more intense emotional states than when piglets are further away from the human. However, it is still unclear whether the changes in grunt structure could also be linked to human-animal communication and more studies are needed to determine it. We did also show that the identity of the human may be of importance, and may generate vocal changes during additional positive contacts that were not associated with changes in behaviour of the human. More systematic studies of human behaviour along with pig behaviour during the human-animal interactions would be needed to have a better understanding of the evolution of HAR, especially interactive interspecific communication as well as providing new procedures to promote positive welfare. We suggest that analysing vocalisations structure may be a good tool to assess the quality of human-pig relationship and help monitor the establishment of a positive HAR.

## Authors contributions

Conceived and designed the experiment (A.S.V., C.T.). Performed the experiment (A.S.V., C.G.). Collection and processing of the acoustic and behavioural data (A.S.V., C.G.). Statistical analyses (A.S.V.). Contributed to the writing of the manuscript (A.S.V., C.T.).

## Acknowledgments

We acknowledge all the technical staff at UEPR: especially Patrick Touanel and Marie-Hélène Lohat, who largely participated in handling the piglets. We thank Eric Siroux who helped building the acoustic chamber at the beginning of the experiment, Remi Resmond for great discussions about statistics and Bliss Elizabeth Bagnato-Conlin for their carefully proof reading of the manuscript. All the authors acknowledge Camille Noûs, affirming the collective and open character of the creation and dissemination of knowledge (Cogitamus Laboratory https://www.cogitamus.fr/indexen.html). This project is part of the SoundWel project in the framework of the Anihwa Eranet and funded by ANR 30001199.

## Data availability

The datasets used for the study are available at (Villain et al., 2022). The folder contains all datasets and a readme to match the type of analysis to the proper dataset. We have made sure to report in the main text of the article which R libraries and which functions in these libraries we used. All formulas of the statistical models are explicit in the text to facilitate transfer of information and replicate the analysis. All libraries are open source as well.

