## Supplementary Tables for "The use of pigs vocalisation structure to assess the quality of human-pig relationship"

### Electronic supplementary material

Table S1 : Anova table of all models computed. The function 'Anova' from the 'car' R package was used to generate p value on full models. Interpretable significant p-values are bolded. When significant interaction were found, post hoc tests were performed (see table S2).

| Fixed effects | Chisq | Df | Pr..Chisq. |
|---|---|---|---|
| **Model #1 : behavioural response Reunion of the Isolation/Reunion test** | | | |
| **ReuPC1** | | | |
| Treatment | 19.077 | 1 | <0.001 |
| Conditioning time | 139.035 | 1 | <0.001 |
| Batch | 42.566 | 1 | <0.001 |
| HumanID | 0.137 | 1 | 0.711 |
| Treatment:Conditioning time | 27.910 | 1 | **<0.001** |
| Treatment:Batch | 0.507 | 1 | 0.476 |
| Treatment:HumanID | 6.009 | 1 | **0.014** |
| Conditioning time:Batch | 20.240 | 1 | **<0.001** |
| Conditioning time:HumanID | 0.646 | 1 | 0.422 |
| **ReuPC2 (sqrt+4)** | | | |
| Treatment | 0.995 | 1 | 0.319 |
| Conditioning time | 3.782 | 1 | 0.052 |
| Batch | 5.118 | 1 | 0.024 |
| HumanID | 1.978 | 1 | 0.160 |
| Treatment:Conditioning time | 0.000 | 1 | 0.989 |
| Treatment:Batch | 0.333 | 1 | 0.564 |
| Treatment:HumanID | 1.752 | 1 | 0.186 |
| Conditioning time:Batch | 14.193 | 1 | **<0.001** |
| Conditioning time:HumanID | 0.189 | 1 | 0.663 |
| **-ReuPC3 (sqrt +3)** | | | |
| Treatment | 6.884 | 1 | 0.009 |
| Conditioning time | 31.456 | 1 | <0.001 |
| Batch | 0.000 | 1 | 0.984 |
| HumanID | 0.385 | 1 | 0.535 |
| Treatment:Conditioning time | 3.658 | 1 | 0.056 |
| Treatment:Batch | 6.966 | 1 | **0.008** |
| Treatment:HumanID | 2.010 | 1 | 0.156 |
| Conditioning time:Batch | 5.445 | 1 | **0.020** |
| Conditioning time:HumanID | 0.247 | 1 | 0.619 |
| **Model #2 : Vocal response Isolation/Reunion tests : Treatment * Phase * Conditioning time** | | | |
| **Call duration (s) (log)** | | | |
| Treatment | 5.503 | 1 | **0.019** |
| Phase | 60.842 | 1 | **<0.001** |
| Conditioning time | 62.883 | 1 | <0.001 |
| HumanID | 0.535 | 1 | 0.465 |

| | | | |
|---|---|---|---|
| Batch | 8.053 | 1 | 0.005 |
| Treatment:Phase | 0.872 | 1 | 0.350 |
| Treatment:Conditioning time | 3.479 | 1 | 0.062 |
| Phase:Conditioning time | 1.894 | 1 | 0.169 |
| Treatment:HumanID | 0.048 | 1 | 0.826 |
| Conditioning time:HumanID | 2.347 | 1 | 0.126 |
| Treatment:Batch | 2.398 | 1 | 0.121 |
| Conditioning time:Batch | 10.844 | 1 | **0.001** |
| Treatment:Phase:Conditioning time | 0.699 | 1 | 0.403 |
| **-VocPC1** | | | |
| Treatment | 0.886 | 1 | 0.346 |
| Phase | 8.501 | 1 | **0.004** |
| Conditioning time | 0.359 | 1 | 0.549 |
| HumanID | 2.519 | 1 | 0.112 |
| Batch | 60.781 | 1 | **<0.001** |
| Treatment:Phase | 0.735 | 1 | 0.391 |
| Treatment:Conditioning time | 0.592 | 1 | 0.442 |
| Phase:Conditioning time | 0.616 | 1 | 0.433 |
| Treatment:HumanID | 0.095 | 1 | 0.758 |
| Conditioning time:HumanID | 0.786 | 1 | 0.375 |
| Treatment:Batch | 0.129 | 1 | 0.720 |
| Conditioning time:Batch | 1.875 | 1 | 0.171 |
| Treatment:Phase:Conditioning time | 0.000 | 1 | 0.995 |
| **VocPC2** | | | |
| Treatment | 0.011 | 1 | 0.918 |
| Phase | 19.116 | 1 | **<0.001** |
| Conditioning time | 245.911 | 1 | <0.001 |
| HumanID | 6.152 | 1 | **0.013** |
| Batch | 2.378 | 1 | 0.123 |
| Treatment:Phase | 3.525 | 1 | 0.060 |
| Treatment:Conditioning time | 0.695 | 1 | 0.405 |
| Phase:Conditioning time | 2.105 | 1 | 0.147 |
| Treatment:HumanID | 0.280 | 1 | 0.597 |
| Conditioning time:HumanID | 0.032 | 1 | 0.858 |
| Treatment:Batch | 0.355 | 1 | 0.552 |
| Conditioning time:Batch | 34.561 | 1 | **<0.001** |
| Treatment:Phase:Conditioning time | 0.314 | 1 | 0.576 |
| **-VocPC3** | | | |
| Treatment | 4.782 | 1 | **0.029** |
| Phase | 6.567 | 1 | **0.010** |
| Conditioning time | 2.945 | 1 | 0.086 |
| HumanID | 0.870 | 1 | 0.351 |
| Batch | 50.730 | 1 | <0.001 |
| Treatment:Phase | 0.721 | 1 | 0.396 |
| Treatment:Conditioning time | 0.102 | 1 | 0.750 |
| Phase:Conditioning time | 2.026 | 1 | 0.155 |
| Treatment:HumanID | 0.087 | 1 | 0.767 |
| Conditioning time:HumanID | 2.002 | 1 | 0.157 |
| Treatment:Batch | 1.905 | 1 | 0.168 |

| | | | |
|---|---|---|---|
| Conditioning time:Batch | 8.468 | 1 | **0.004** |
| Treatment:Phase:Conditioning time | 0.624 | 1 | 0.429 |

| **Model #3 : Vocal response Reunion of the Isolation/Reunion tests : conditioning time * Treatment * In prox. area** | | | |
|---|---|---|---|
| **Call duration (s) (log)** | | | |
| Conditioning time | 37.742 | 1 | <0.001 |
| Treatment | 0.943 | 1 | 0.331 |
| In prox. area | 48.590 | 1 | <0.001 |
| HumanID | 2.208 | 1 | 0.137 |
| Batch | 4.987 | 1 | 0.026 |
| Conditioning time:Treatment | 1.892 | 1 | 0.169 |
| Conditioning time:In prox. area | 4.913 | 1 | 0.027 |
| Treatment:In prox. area | 16.021 | 1 | <0.001 |
| Conditioning time:HumanID | 0.526 | 1 | 0.468 |
| Conditioning time:Batch | 29.430 | 1 | **<0.001** |
| Treatment:Batch | 0.172 | 1 | 0.678 |
| Treatment:HumanID | 0.004 | 1 | 0.947 |
| In prox. area:HumanID | 2.058 | 1 | 0.151 |
| In prox. area:Batch | 0.010 | 1 | 0.919 |
| Conditioning time:Treatment:In prox. area | 6.541 | 1 | **0.011** |
| **-VocPC1** | | | |
| Conditioning time | 0.391 | 1 | 0.532 |
| Treatment | 0.026 | 1 | 0.873 |
| In prox. area | 0.973 | 1 | 0.324 |
| HumanID | 3.006 | 1 | 0.083 |
| Batch | 36.673 | 1 | **<0.001** |
| Conditioning time:Treatment | 0.802 | 1 | 0.371 |
| Conditioning time:In prox. area | 0.600 | 1 | 0.439 |
| Treatment:In prox. area | 14.375 | 1 | <0.001 |
| Conditioning time:HumanID | 0.261 | 1 | 0.609 |
| Conditioning time:Batch | 3.911 | 1 | 0.048 |
| Treatment:Batch | 1.671 | 1 | 0.196 |
| Treatment:HumanID | 0.272 | 1 | 0.602 |
| In prox. area:HumanID | 2.024 | 1 | 0.155 |
| In prox. area:Batch | 2.939 | 1 | 0.086 |
| Conditioning time:Treatment:In prox. area | 4.987 | 1 | **0.026** |
| **VocPC2** | | | |
| Conditioning time | 110.726 | 1 | <0.001 |
| Treatment | 0.351 | 1 | 0.554 |
| In prox. area | 26.883 | 1 | <0.001 |
| HumanID | 4.938 | 1 | **0.026** |
| Batch | 2.275 | 1 | 0.131 |
| Conditioning time:Treatment | 0.016 | 1 | 0.900 |
| Conditioning time:In prox. area | 10.339 | 1 | **0.001** |
| Treatment:In prox. area | 4.225 | 1 | **0.040** |
| Conditioning time:HumanID | 0.037 | 1 | 0.848 |
| Conditioning time:Batch | 37.624 | 1 | **<0.001** |
| Treatment:Batch | 0.342 | 1 | 0.559 |

| | | | |
|---|---|---|---|
| Treatment:HumanID | 0.403 | 1 | 0.525 |
| In prox. area:HumanID | 0.020 | 1 | 0.887 |
| In prox. area:Batch | 8.818 | 1 | **0.003** |
| Conditioning time:Treatment:In prox. area | 3.353 | 1 | 0.067 |
| **-VocPC3** | | | |
| Conditioning time | 6.221 | 1 | 0.013 |
| Treatment | 3.158 | 1 | 0.076 |
| In prox. area | 8.537 | 1 | 0.003 |
| HumanID | 1.180 | 1 | 0.277 |
| Batch | 40.179 | 1 | <0.001 |
| Conditioning time:Treatment | 0.371 | 1 | 0.542 |
| Conditioning time:In prox. area | 3.245 | 1 | 0.072 |
| Treatment:In prox. area | 1.308 | 1 | 0.253 |
| Conditioning time:HumanID | 0.154 | 1 | 0.695 |
| Conditioning time:Batch | 8.632 | 1 | **0.003** |
| Treatment:Batch | 2.241 | 1 | 0.134 |
| Treatment:HumanID | 0.046 | 1 | 0.830 |
| In prox. area:HumanID | 2.982 | 1 | 0.084 |
| In prox. area:Batch | 10.363 | 1 | **0.001** |
| Conditioning time:Treatment:In prox. area | 4.893 | 1 | **0.027** |
| **Model #4 : Behavioural response during sessions of conditioning** | | | |
| **CondPC1** | | | |
| Trial number | 59.317 | 1 | **<0.001** |
| Treatment | 0.128 | 1 | 0.721 |
| HumanID | 1.320 | 1 | 0.251 |
| Batch | 14.497 | 1 | **<0.001** |
| Trial number:Treatment | 2.545 | 1 | 0.111 |
| Trial number:HumanID | 0.023 | 1 | 0.880 |
| Trial number:Batch | 0.626 | 1 | 0.429 |
| Treatment:Batch | 1.663 | 1 | 0.197 |
| Treatment:HumanID | 0.437 | 1 | 0.508 |
| **CondPC2** | | | |
| Trial number | 48.618 | 1 | **<0.001** |
| Treatment | 12.806 | 1 | **<0.001** |
| HumanID | 0.226 | 1 | 0.635 |
| Batch | 10.056 | 1 | **0.002** |
| Trial number:Treatment | 0.041 | 1 | 0.839 |
| Trial number:HumanID | 0.000 | 1 | 0.999 |
| Trial number:Batch | 0.085 | 1 | 0.771 |
| Treatment:Batch | 2.007 | 1 | 0.157 |
| Treatment:HumanID | 0.907 | 1 | 0.341 |
| **CondPC3** | | | |
| Trial number | 0.006 | 1 | 0.939 |
| Treatment | 6.969 | 1 | **0.008** |
| HumanID | 0.375 | 1 | 0.541 |
| Batch | 0.015 | 1 | 0.903 |
| Trial number:Treatment | 0.616 | 1 | 0.432 |
| Trial number:HumanID | 0.109 | 1 | 0.741 |

| | | | |
|---|---|---|---|
| Trial number:Batch | 0.166 | 1 | 0.684 |
| Treatment:Batch | 0.078 | 1 | 0.780 |
| Treatment:HumanID | 0.143 | 1 | 0.705 |
| **Missed contact attempts by Human ID (binomial)** | | | |
| Trial number | 23.159 | 1 | <0.001 |
| Treatment | 9.563 | 1 | **0.002** |
| HumanID | 0.463 | 1 | 0.496 |
| Batch | 12.833 | 1 | <0.001 |
| Trial number:Treatment | 0.218 | 1 | 0.640 |
| Trial number:HumanID | 0.058 | 1 | 0.809 |
| Trial number:Batch | 4.485 | 1 | **0.034** |
| Treatment:Batch | 1.274 | 1 | 0.259 |
| Treatment:HumanID | 1.073 | 1 | 0.300 |
| **Model #5 : Vocal response during session of conditioning : Trial number * Treatment * In prox. area** | | | |
| **Call duration (s) (log)** | | | |
| Trial number | 48.880 | 1 | <0.001 |
| Treatment | 5.192 | 1 | 0.023 |
| In prox. area | 160.565 | 1 | <0.001 |
| HumanID | 0.090 | 1 | 0.765 |
| Batch | 11.814 | 1 | 0.001 |
| Trial number:Treatment | 0.384 | 1 | 0.536 |
| Trial number:In prox. area | 0.584 | 1 | 0.445 |
| Treatment:In prox. area | 15.779 | 1 | **<0.001** |
| Trial number:HumanID | 1.252 | 1 | 0.263 |
| Trial number:Batch | 5.374 | 1 | **0.020** |
| Treatment:Batch | 6.716 | 1 | **0.010** |
| Treatment:HumanID | 0.712 | 1 | 0.399 |
| In prox. area:HumanID | 0.004 | 1 | 0.951 |
| In prox. area:Batch | 0.105 | 1 | 0.746 |
| Trial number:Treatment:In prox. area | 0.019 | 1 | 0.889 |
| **-VocPC1** | | | |
| Trial number | 12.233 | 1 | <0.001 |
| Treatment | 0.043 | 1 | 0.835 |
| In prox. area | 2.225 | 1 | 0.136 |
| HumanID | 4.696 | 1 | **0.030** |
| Batch | 62.339 | 1 | <0.001 |
| Trial number:Treatment | 1.091 | 1 | 0.296 |
| Trial number:In prox. area | 3.968 | 1 | **0.046** |
| Treatment:In prox. area | 1.089 | 1 | 0.297 |
| Trial number:HumanID | 0.155 | 1 | 0.694 |
| Trial number:Batch | 0.099 | 1 | 0.753 |
| Treatment:Batch | 6.990 | 1 | **0.008** |
| Treatment:HumanID | 0.606 | 1 | 0.436 |
| In prox. area:HumanID | 2.204 | 1 | 0.138 |
| In prox. area:Batch | 5.703 | 1 | **0.017** |
| Trial number:Treatment:In prox. area | 0.275 | 1 | 0.600 |
| **VocPC2** | | | |
| Trial number | 85.956 | 1 | <0.001 |

| | | | |
|---|---|---|---|
| Treatment | 0.217 | 1 | 0.642 |
| In prox. area | 13.440 | 1 | <0.001 |
| HumanID | 2.932 | 1 | 0.087 |
| Batch | 6.712 | 1 | 0.010 |
| Trial number:Treatment | 0.507 | 1 | 0.477 |
| Trial number:In prox. area | 6.107 | 1 | **0.013** |
| Treatment:In prox. area | 7.622 | 1 | **0.006** |
| Trial number:HumanID | 0.016 | 1 | 0.899 |
| Trial number:Batch | 8.261 | 1 | **0.004** |
| Treatment:Batch | 1.482 | 1 | 0.223 |
| Treatment:HumanID | 2.318 | 1 | 0.128 |
| In prox. area:HumanID | 11.895 | 1 | **0.001** |
| In prox. area:Batch | 1.792 | 1 | 0.181 |
| Trial number:Treatment:In prox. area | 0.049 | 1 | 0.824 |
| **VocPC3** | | | |
| Trial number | 14.564 | 1 | <0.001 |
| Treatment | 2.710 | 1 | 0.100 |
| In prox. area | 4.578 | 1 | 0.032 |
| HumanID | 0.652 | 1 | 0.419 |
| Batch | 44.701 | 1 | **<0.001** |
| Trial number:Treatment | 2.485 | 1 | 0.115 |
| Trial number:In prox. area | 0.315 | 1 | 0.575 |
| Treatment:In prox. area | 2.502 | 1 | 0.114 |
| Trial number:HumanID | 7.978 | 1 | **0.005** |
| Trial number:Batch | 0.456 | 1 | 0.499 |
| Treatment:Batch | 0.029 | 1 | 0.865 |
| Treatment:HumanID | 0.000 | 1 | 0.984 |
| In prox. area:HumanID | 1.696 | 1 | 0.193 |
| In prox. area:Batch | 0.001 | 1 | 0.970 |
| Trial number:Treatment:In prox. area | 0.183 | 1 | 0.669 |

Table S2 : Table of contrasts from posthoc tests following significant interaction using the Anova on the model, pairwise comparison with Tukey correction. P-values were generated using the ‘emmeans’ (categorical fixed effect) and ‘lstrends’ (continuous fixed effect) functions of the ‘Emmeans’ R package. Estimates indicated are either between groups or slope comparisons, depending on the categorial or continuous variable (trial number). When fixed effect interacting with the batch, the batch number was fixed to compare the fixed effect within each batch. When three way interaction were significant, one factor was fixed to compare the interacting effect of the two other (conditioning time was fixed).

| **contrast** | **fixed comparison factor if any** | **estimate** | **SE** | **ratio** | **p.value** |
|---|---|---|---|---|---|
| **Model #1 : behavioural response of the Reunion of Isolation/Reunion test** | | | | | |
| **ReuPC1** | | | | | |
| Treatment * Conditioning time | | | | | |
| H after - H+ after | - | 0.022 | 0.311 | 0.072 | 1.000 |
| H after - H before | - | 3.430 | 0.284 | 12.073 | <0.001 |
| H after - H+ before | - | 1.330 | 0.311 | 4.271 | <0.001 |
| H+ after - H before | - | 3.407 | 0.311 | 10.945 | <0.001 |

| | | | | | |
|---|---|---|---|---|---|
| H+ after - H+ before | | 1.307 | 0.284 | 4.602 | |
| H before – H+ before | - | -2.100 | 0.311 | -6.746 | <0.001 |
| Conditioning time \| Batch | | | | | |
| after - before | 1 | 1.446 | 0.287 | 5.039 | <0.001 |
| after - before | 2 | 3.291 | 0.287 | 11.466 | <0.001 |
| Treatment * HumanID | | | | | |
| H AH - H+ AH | | -1.634 | 0.340 | -4.808 | <0.001 |
| H AH - H AV | | -0.685 | 0.343 | -1.995 | 0.202 |
| H AH - H+ AV | | -1.129 | 0.340 | -3.321 | 0.009 |
| H+ AH - H AV | | 0.949 | 0.340 | 2.792 | 0.035 |
| H+ AH - H+ AV | | 0.505 | 0.343 | 1.471 | 0.462 |
| H AV - H+ AV | | -0.444 | 0.340 | -1.306 | 0.563 |
| **ReuPC2 (sqrt+4)** | | | | | |
| Conditioning time \| Batch | | | | | |
| after - before | 1 | 0.137 | 0.103 | 1.330 | 0.189 |
| after - before | 2 | -0.416 | 0.103 | -4.052 | <0.001 |
| **-ReuPC3 (sqrt +3)** | | | | | |
| Treatment * Conditioning time | | | | | |
| H after - H+ after | - | -0.252 | 0.078 | -3.237 | 0.009 |
| H after - H before | - | 0.187 | 0.072 | 2.613 | 0.054 |
| H after - H+ before | - | 0.129 | 0.078 | 1.657 | 0.352 |
| H+ after - H before | - | 0.439 | 0.078 | 5.642 | <0.001 |
| H+ after - H+ before | - | 0.381 | 0.072 | 5.318 | <0.001 |
| H before - H+ before | - | -0.058 | 0.078 | -0.748 | 0.877 |
| Treatment \| Batch | | | | | |
| H - H+ | 1 | -0.314 | 0.084 | -3.721 | <0.001 |
| H - H+ | 2 | 0.004 | 0.084 | 0.049 | 0.961 |
| Conditioning time \| Batch | | | | | |
| after - before | 1 | 0.404 | 0.072 | 5.592 | <0.001 |
| after - before | 2 | 0.163 | 0.072 | 2.258 | 0.028 |
| **Model #2 : Vocal response Isolation/Reunion tests : Treatment * Phase * Conditioning time** | | | | | |
| **Call duration (s) (log)** | | | | | |
| Conditioning time \| Batch | | | | | |
| after - before | 1 | 0.171 | 0.045 | 3.760 | <0.001 |
| after - before | 2 | 0.398 | 0.052 | 7.680 | <0.001 |
| **VocPC2** | | | | | |
| Conditioning time \| Batch | | | | | |
| after - before | 1 | -0.832 | 0.101 | -8.232 | <0.001 |
| after - before | 2 | -1.755 | 0.120 | -14.595 | <0.001 |
| **-VocPC3** | | | | | |
| Conditioning time \| Batch | | | | | |
| after - before | 1 | 0.308 | 0.100 | 3.085 | 0.002 |
| after - before | 2 | -0.136 | 0.115 | -1.181 | 0.238 |
| **Model #3 : Vocal response during the Reunion of Isolation/Reunion tests : conditioning time * Treatment * In prox. area** | | | | | |

| | | | | | |
|---|---|---|---|---|---|
| **Call duration (s) (log)** | | | | | |
| Treatment * In prox. area \| conditioning time | | | | | |
| H 0 - H+ 0 | after | 0.040 | 0.071 | 0.564 | 0.943 |
| H 0 - H 1 | after | 0.123 | 0.030 | 4.097 | <0.001 |
| H 0 - H+ 1 | after | 0.090 | 0.073 | 1.224 | 0.612 |
| H+ 0 - H 1 | after | 0.083 | 0.074 | 1.115 | 0.680 |
| H+ 0 - H+ 1 | after | 0.050 | 0.025 | 1.989 | 0.192 |
| H 1 - H+ 1 | after | -0.033 | 0.076 | -0.433 | 0.973 |
| H 0 - H+ 0 | before | 0.187 | 0.079 | 2.384 | 0.080 |
| H 0 - H 1 | before | 0.312 | 0.049 | 6.329 | <0.001 |
| H 0 - H+ 1 | before | 0.254 | 0.080 | 3.185 | 0.008 |
| H+ 0 - H 1 | before | 0.124 | 0.088 | 1.418 | 0.488 |
| H+ 0 - H+ 1 | before | 0.066 | 0.030 | 2.186 | 0.127 |
| H 1 - H+ 1 | before | -0.058 | 0.087 | -0.664 | 0.911 |
| Conditioning time \| Batch | | | | | |
| after - before | 1 | 0.084 | 0.049 | 1.734 | 0.083 |
| after - before | 2 | 0.492 | 0.059 | 8.334 | <0.001 |
| **-VocPC1** | | | | | |
| Treatment * In prox. area \| conditioning time | | | | | |
| H 0 - H+ 0 | after | -0.345 | 0.416 | -0.829 | 0.841 |
| H 0 - H 1 | after | -0.227 | 0.184 | -1.232 | 0.607 |
| H 0 - H+ 1 | after | -0.124 | 0.431 | -0.288 | 0.992 |
| H+ 0 - H 1 | after | 0.118 | 0.436 | 0.271 | 0.993 |
| H+ 0 - H+ 1 | after | 0.221 | 0.153 | 1.443 | 0.473 |
| H 1 - H+ 1 | after | 0.103 | 0.450 | 0.228 | 0.996 |
| H 0 - H+ 0 | before | -0.052 | 0.473 | -0.109 | 1.000 |
| H 0 - H 1 | before | -1.014 | 0.304 | -3.340 | 0.005 |
| H 0 - H+ 1 | before | 0.311 | 0.480 | 0.647 | 0.917 |
| H+ 0 - H 1 | before | -0.962 | 0.531 | -1.811 | 0.268 |
| H+ 0 - H+ 1 | before | 0.362 | 0.187 | 1.939 | 0.212 |
| H 1 - H+ 1 | before | 1.324 | 0.530 | 2.500 | 0.060 |
| Conditioning time \| Batch | | | | | |
| after - before | 1 | 0.552 | 0.395 | 1.397 | 0.162 |
| after - before | 2 | -0.635 | 0.458 | -1.387 | 0.166 |
| **VocPC2*** | | | | | |
| *Note : due to a three way interaction close to significance level, contrasts were generated with the three way interaction and with the two ways interactions of interests | | | | | |
| Treatment * In prox. area \| conditioning time | | | | | |
| H 0 - H+ 0 | after | 0.112 | 0.215 | 0.519 | 0.955 |
| H 0 - H 1 | after | -0.216 | 0.093 | -2.326 | 0.092 |
| H 0 - H+ 1 | after | -0.022 | 0.223 | -0.100 | 1.000 |
| H+ 0 - H 1 | after | -0.328 | 0.225 | -1.454 | 0.466 |
| H+ 0 - H+ 1 | after | -0.134 | 0.077 | -1.733 | 0.306 |
| H 1 - H+ 1 | after | 0.193 | 0.232 | 0.834 | 0.838 |
| H 0 - H+ 0 | before | -0.016 | 0.238 | -0.067 | 1.000 |
| H 0 - H 1 | before | -0.798 | 0.152 | -5.239 | <0.001 |
| H 0 - H+ 1 | before | -0.351 | 0.241 | -1.454 | 0.466 |

| | | | | | |
|---|---|---|---|---|---|
| H+ 0 - H 1 | before | -0.783 | 0.267 | -2.933 | 0.018 |
| H+ 0 - H+ 1 | before | -0.335 | 0.094 | -3.561 | 0.002 |
| H 1 - H+ 1 | before | 0.448 | 0.266 | 1.685 | 0.332 |
| In prox. area \| Conditioning time | | | | | |
| 0 - 1 | after | -0.175 | 0.061 | -2.876 | 0.004 |
| 0 - 1 | before | -0.567 | 0.093 | -6.124 | <0.001 |
| Treatment * In prox. area | | | | | |
| H 0 - H+ 0 | | 0.048 | 0.198 | 0.242 | 0.995 |
| H 0 - H 1 | | -0.507 | 0.092 | -5.539 | <0.001 |
| H 0 - H+ 1 | | -0.187 | 0.202 | -0.926 | 0.791 |
| H+ 0 - H 1 | | -0.555 | 0.211 | -2.634 | 0.042 |
| H+ 0 - H+ 1 | | -0.235 | 0.061 | -3.827 | 0.001 |
| H 1 - H+ 1 | | 0.320 | 0.212 | 1.513 | 0.429 |
| In prox. area \| Batch | | | | | |
| 0 - 1 | 1 | -0.223 | 0.064 | -3.480 | 0.001 |
| 0 - 1 | 2 | -0.519 | 0.086 | -6.051 | <0.001 |
| Conditioning time \| Batch | | | | | |
| after - before | 1 | -0.680 | 0.140 | -4.864 | <0.001 |
| after - before | 2 | -2.009 | 0.172 | -11.710 | <0.001 |
| **-VocPC3** | | | | | |
| Treatment * In prox. area | | | | | |
| H 0 - H+ 0 | after | 0.227 | 0.132 | 1.726 | 0.310 |
| H 0 - H 1 | after | 0.125 | 0.073 | 1.703 | 0.322 |
| H 0 - H+ 1 | after | 0.324 | 0.139 | 2.332 | 0.091 |
| H+ 0 - H 1 | after | -0.102 | 0.142 | -0.722 | 0.888 |
| H+ 0 - H+ 1 | after | 0.097 | 0.061 | 1.580 | 0.390 |
| H 1 - H+ 1 | after | 0.199 | 0.148 | 1.345 | 0.534 |
| H 0 - H+ 0 | before | 0.033 | 0.153 | 0.218 | 0.996 |
| H 0 - H 1 | before | -0.266 | 0.120 | -2.207 | 0.121 |
| H 0 - H+ 1 | before | 0.105 | 0.156 | 0.674 | 0.907 |
| H+ 0 - H 1 | before | -0.299 | 0.179 | -1.665 | 0.342 |
| H+ 0 - H+ 1 | before | 0.072 | 0.074 | 0.961 | 0.772 |
| H 1 - H+ 1 | before | 0.370 | 0.178 | 2.077 | 0.161 |
| Conditioning time \| Batch | | | | | |
| after - before | 1 | 0.334 | 0.103 | 3.237 | 0.001 |
| after - before | 2 | -0.136 | 0.127 | -1.071 | 0.284 |
| **Model #4 : Behavioural response during session of the conditioning** | | | | | |
| **Occurence of missed contact from human** | | | | | |
| Treatment | | | | | |
| H - H+ | 0.812 | 0.271 - | | 3.003 | 0.003 |
| Trial number \| Batch | | | | | |
| 1 - 2 | -0.177 | 0.083 - | | -2.118 | 0.034 |
| **Model #5 : Vocal response during session of the conditioning Trial number * Treatment * In prox. area** | | | | | |
| **Call duration (s) (log)** | | | | | |

| | | | | | |
|---|---|---|---|---|---|
| Treatment * In prox. area | | | | | |
| H 0 - H+ 0 | - | 0.121 | 0.053 | 2.293 | 0.100 |
| H 0 - H 1 | - | 0.357 | 0.035 | 10.168 | <0.001 |
| H 0 - H+ 1 | - | 0.304 | 0.058 | 5.220 | <0.001 |
| H+ 0 - H 1 | - | 0.236 | 0.062 | 3.797 | <0.001 |
| H+ 0 - H+ 1 | - | 0.182 | 0.027 | 6.862 | <0.001 |
| H 1 - H+ 1 | - | -0.053 | 0.067 | -0.801 | 0.854 |
| Trial number \| Batch | | | | | |
| 1 - 2 | - | -0.028 | 0.012 | -2.318 | 0.020 |
| Treatment \| Batch | | | | | |
| H - H+ | 1 | -0.097 | 0.074 | -1.307 | 0.191 |
| H - H+ | 2 | 0.165 | 0.077 | 2.155 | 0.031 |
| **-VocPC1** | | | | | |
| Trial number * In prox. area | | | | | |
| 0 - 1 | - | -0.050 | 0.028 | -1.804 | 0.071 |
| Treatment \| Batch | | | | | |
| H - H+ | 1 | 0.617 | 0.312 | 1.977 | 0.048 |
| H - H+ | 2 | -0.537 | 0.321 | -1.675 | 0.094 |
| In prox. area \| Batch | | | | | |
| 0 - 1 | 1 | -0.291 | 0.113 | -2.568 | 0.010 |
| 0 - 1 | 2 | 0.184 | 0.154 | 1.191 | 0.234 |
| **VocPC2** | | | | | |
| Treatment * In prox. area | | | | | |
| H 0 - H+ 0 | - | -0.097 | 0.149 | -0.649 | 0.916 |
| H 0 - H 1 | - | -0.401 | 0.081 | -4.930 | <0.001 |
| H 0 - H+ 1 | - | -0.216 | 0.159 | -1.356 | 0.527 |
| H+ 0 - H 1 | - | -0.304 | 0.167 | -1.821 | 0.263 |
| H+ 0 - H+ 1 | - | -0.119 | 0.062 | -1.936 | 0.213 |
| H 1 - H+ 1 | - | 0.185 | 0.176 | 1.049 | 0.720 |
| Trial number * In prox. area | | | | | |
| 0 - 1 | - | -0.036 | 0.016 | -2.343 | 0.019 |
| Trial number \| Batch | | | | | |
| 1 - 2 | - | 0.056 | 0.019 | 2.874 | 0.004 |
| In prox. area * HumanID | | | | | |
| 0 AH - 1 AH | - | -0.085 | 0.068 | -1.245 | 0.598 |
| 0 AH - 0 AV | - | 0.290 | 0.151 | 1.919 | 0.220 |
| 0 AH - 1 AV | - | -0.145 | 0.167 | -0.870 | 0.820 |
| 1 AH - 0 AV | - | 0.375 | 0.164 | 2.293 | 0.100 |
| 1 AH - 1 AV | - | -0.060 | 0.178 | -0.336 | 0.987 |
| 0 AV - 1 AV | - | -0.435 | 0.075 | -5.769 | <0.001 |
| **VocPC3** | | | | | |
| Trial number * HumanID | | | | | |
| AH - AV | - | -0.040 | 0.014 | -2.824 | 0.005 |

Table S3 : Table of estimates (standard error SE and 95% confidence intervals) from from models, computed using the ‘emmeans’ (categorical fixed effect) and ‘lstrends’ (continuous fixed effect) functions of the ‘Emmeans’ R package. Estimates indicated are either for groups or slopes, depending on the categorial or continuous variable (trial number). Estimates for all interpretable fixed effect, interacting or not and significant or nor are indicated.

| factor | first interaction (if any) | second interaction (if any) | estimate | SE | Lower 95%confidence int. | Upper 95%confidence int. |
|---|---|---|---|---|---|---|

| Model #1 : behavioural response during the Reunion of Isolation/Reunion test | | | | | |
|---|---|---|---|---|---|
| **ReuPC1** | | | | | |
| Treatment * Conditioning time | | | | | |
| H | after | 1.195 | 0.220 | 0.759 | 1.632 |
| H+ | after | 1.173 | 0.220 | 0.737 | 1.610 |
| H | before | -2.234 | 0.220 | -2.671 | -1.798 |
| H+ | before | -0.134 | 0.220 | -0.571 | 0.302 |
| Conditioning time \| Batch | | | | | |
| after | 1 | 1.515 | 0.222 | 1.074 | 1.956 |
| before | 1 | 0.069 | 0.222 | -0.372 | 0.510 |
| after | 2 | 0.854 | 0.222 | 0.413 | 1.295 |
| before | 2 | -2.437 | 0.222 | -2.878 | -1.996 |
| Treatment * HumanID | | | | | |
| H | AH | -0.862 | 0.240 | -1.344 | -0.380 |
| H+ | AH | 0.772 | 0.240 | 0.290 | 1.254 |
| H | AV | -0.177 | 0.240 | -0.659 | 0.305 |
| H+ | AV | 0.267 | 0.240 | -0.215 | 0.749 |
| **ReuPC2 (sqrt+4)** | | | | | |
| Conditioning time \| Batch | | | | | |
| after | 1 | 2.106 | 0.073 | 1.962 | 2.250 |
| before | 1 | 1.970 | 0.073 | 1.826 | 2.114 |
| after | 2 | 1.664 | 0.073 | 1.520 | 1.808 |
| before | 2 | 2.080 | 0.073 | 1.936 | 2.224 |
| Treatment | | | | | |
| H | | 1.991 | 0.051 | 1.889 | 2.093 |
| H+ | | 1.919 | 0.051 | 1.817 | 2.021 |
| HumanID | | | | | |
| AH | | 1.903 | 0.051 | 1.800 | 2.006 |
| AV | | 2.007 | 0.051 | 1.904 | 2.110 |
| **-ReuPC3 (sqrt +3)** | | | | | |
| Treatment * Conditioning time | | | | | |
| H | after | 1.712 | 0.055 | 1.603 | 1.821 |
| H+ | after | 1.964 | 0.055 | 1.855 | 2.073 |
| H | before | 1.525 | 0.055 | 1.416 | 1.634 |
| H+ | before | 1.583 | 0.055 | 1.474 | 1.692 |
| Treatment \| Batch | | | | | |
| H | 1 | 1.539 | 0.060 | 1.419 | 1.658 |
| H+ | 1 | 1.853 | 0.060 | 1.733 | 1.972 |
| H | 2 | 1.699 | 0.060 | 1.579 | 1.819 |
| H+ | 2 | 1.695 | 0.060 | 1.575 | 1.814 |

| | | | | | |
|---|---|---|---|---|---|
| Conditioning time \| Batch | | | | | |
| after | 1 | 1.898 | 0.056 | 1.788 | 2.008 |
| before | 1 | 1.494 | 0.056 | 1.383 | 1.604 |
| after | 2 | 1.779 | 0.056 | 1.668 | 1.889 |
| before | 2 | 1.615 | 0.056 | 1.505 | 1.725 |
| HumanID | | | | | |
| AH | | 1.678 | 0.042 | 1.593 | 1.762 |
| AV | | 1.715 | 0.042 | 1.630 | 1.800 |
| **Model #2 : Vocal response Isolation/Reunion tests : Treatment * Phase * Conditioning time** | | | | | |
| **Call duration (s) (log)** | | | | | |
| Conditioning time \| Batch | | | | | |
| after | 1 | -1.166 | 0.039 | -1.243 | -1.090 |
| before | 1 | -1.337 | 0.041 | -1.418 | -1.257 |
| after | 2 | -0.928 | 0.041 | -1.008 | -0.848 |
| before | 2 | -1.326 | 0.048 | -1.420 | -1.233 |
| Treatment | | | | | |
| H | | -1.125 | 0.035 | -1.194 | -1.056 |
| H+ | | -1.254 | 0.033 | -1.320 | -1.189 |
| Phase | | | | | |
| isolation | | -1.063 | 0.030 | -1.122 | -1.003 |
| reunion H | | -1.316 | 0.029 | -1.373 | -1.260 |
| HumanID | | | | | |
| AH | | -1.204 | 0.034 | -1.271 | -1.137 |
| AV | | -1.175 | 0.035 | -1.243 | -1.107 |
| **-VocPC1** | | | | | |
| Treatment | | | | | |
| H | | 0.436 | 0.197 | 0.050 | 0.821 |
| H+ | | 0.687 | 0.186 | 0.323 | 1.051 |
| Phase | | | | | |
| isolation | | 0.341 | 0.161 | 0.025 | 0.656 |
| reunion H | | 0.782 | 0.154 | 0.480 | 1.083 |
| HumanID | | | | | |
| AH | | 0.763 | 0.190 | 0.391 | 1.136 |
| AV | | 0.359 | 0.193 | -0.020 | 0.738 |
| Batch | | | | | |
| 1 | | 1.605 | 0.183 | 1.247 | 1.963 |
| 2 | | -0.482 | 0.202 | -0.877 | -0.087 |
| Conditioning time | | | | | |
| after | | 0.594 | 0.167 | 0.268 | 0.921 |
| before | | 0.528 | 0.185 | 0.166 | 0.890 |
| **VocPC2** | | | | | |
| Conditioning time \| Batch | | | | | |
| after | 1 | -0.645 | 0.129 | -0.898 | -0.392 |
| before | 1 | 0.187 | 0.133 | -0.073 | 0.448 |

| | | | | | | |
|---|---|---|---|---|---|---|
| after | 2 | | -1.290 | 0.137 | -1.558 | -1.022 |
| before | 2 | | 0.465 | 0.150 | 0.172 | 0.759 |
| Treatment | | | | | | |
| H | | | -0.340 | 0.128 | -0.591 | -0.089 |
| H+ | | | -0.301 | 0.121 | -0.539 | -0.064 |
| Phase | | | | | | |
| isolation | | | -0.464 | 0.096 | -0.653 | -0.276 |
| reunion H | | | -0.177 | 0.094 | -0.361 | 0.007 |
| HumanID | | | | | | |
| AH | | | -0.097 | 0.125 | -0.342 | 0.147 |
| AV | | | -0.544 | 0.126 | -0.790 | -0.297 |
| **-VocPC3** | | | | | | |
| Conditioning time \| Batch | | | | | | |
| after | 1 | | -0.415 | 0.086 | -0.583 | -0.248 |
| before | 1 | | -0.724 | 0.089 | -0.898 | -0.549 |
| after | 2 | | 0.142 | 0.091 | -0.036 | 0.319 |
| before | 2 | | 0.277 | 0.104 | 0.074 | 0.481 |
| Treatment | | | | | | |
| H | | | -0.068 | 0.077 | -0.219 | 0.083 |
| H+ | | | -0.292 | 0.073 | -0.434 | -0.149 |
| Phase | | | | | | |
| isolation | | | -0.107 | 0.060 | -0.225 | 0.010 |
| reunion H | | | -0.253 | 0.058 | -0.366 | -0.140 |
| HumanID | | | | | | |
| AH | | | -0.136 | 0.074 | -0.282 | 0.010 |
| AV | | | -0.224 | 0.076 | -0.372 | -0.076 |
| **Model #3 : Vocal response during the Reunion of Isolation/Reunion tests : conditioning time * Treatment * In prox. area** | | | | | | |
| **Call duration (s) (log)** | | | | | | |
| Treatment * In prox. area | | | | | | |
| H | 0 | after | -1.154 | 0.052 | -1.256 | -1.053 |
| H+ | 0 | after | -1.194 | 0.049 | -1.289 | -1.099 |
| H | 1 | after | -1.277 | 0.056 | -1.386 | -1.167 |
| H+ | 1 | after | -1.244 | 0.052 | -1.345 | -1.142 |
| H | 0 | before | -1.317 | 0.059 | -1.434 | -1.201 |
| H+ | 0 | before | -1.505 | 0.052 | -1.607 | -1.402 |
| H | 1 | before | -1.629 | 0.071 | -1.768 | -1.490 |
| H+ | 1 | before | -1.571 | 0.054 | -1.677 | -1.465 |
| Conditioning time \| Batch | | | | | | |
| after | 1 | | -1.373 | 0.050 | -1.471 | -1.276 |
| before | 1 | | -1.458 | 0.050 | -1.556 | -1.360 |
| after | 2 | | -1.061 | 0.051 | -1.162 | -0.961 |
| before | 2 | | -1.553 | 0.062 | -1.674 | -1.433 |
| HumanID | | | | | | |
| AH | | | -1.412 | 0.045 | -1.501 | -1.324 |

| | | | | | | |
|---|---|---|---|---|---|---|
| AV | | | -1.311 | 0.046 | -1.402 | -1.220 |
| **-VocPC1** | | | | | | |
| Treatment * In prox. area \| conditioning time | | | | | | |
| H | 0 | after | 0.718 | 0.303 | 0.124 | 1.312 |
| H+ | 0 | after | 1.063 | 0.285 | 0.505 | 1.622 |
| H | 1 | after | 0.945 | 0.330 | 0.298 | 1.592 |
| H+ | 1 | after | 0.842 | 0.306 | 0.242 | 1.442 |
| H | 0 | before | 0.745 | 0.360 | 0.039 | 1.450 |
| H+ | 0 | before | 0.796 | 0.312 | 0.185 | 1.408 |
| H | 1 | before | 1.758 | 0.433 | 0.911 | 2.606 |
| H+ | 1 | before | 0.434 | 0.322 | -0.198 | 1.066 |
| Conditioning time \| Batch | | | | | | |
| after | 1 | | 2.072 | 0.293 | 1.498 | 2.646 |
| before | 1 | | 1.520 | 0.294 | 0.944 | 2.096 |
| after | 2 | | -0.288 | 0.300 | -0.875 | 0.299 |
| before | 2 | | 0.347 | 0.375 | -0.389 | 1.082 |
| HumanID | | | | | | |
| AH | | | 1.145 | 0.228 | 0.699 | 1.592 |
| AV | | | 0.680 | 0.237 | 0.216 | 1.144 |
| **VocPC2*** | | | | | | |
| *Note : due to a three way interaction close to significance level, contrasts were generating with the three way interaction or with the two ways interactions of interests | | | | | | |
| Treatment * In prox. area \| conditioning time | | | | | | |
| H | 0 | after | -0.773 | 0.157 | -1.081 | -0.465 |
| H+ | 0 | after | -0.884 | 0.147 | -1.173 | -0.596 |
| H | 1 | after | -0.557 | 0.170 | -0.891 | -0.223 |
| H+ | 1 | after | -0.750 | 0.158 | -1.059 | -0.441 |
| H | 0 | before | 0.312 | 0.180 | -0.040 | 0.664 |
| H+ | 0 | before | 0.328 | 0.159 | 0.016 | 0.640 |
| H | 1 | before | 1.110 | 0.216 | 0.687 | 1.534 |
| H+ | 1 | before | 0.663 | 0.164 | 0.342 | 0.984 |
| In prox. area \| Conditioning time | | | | | | |
| 0 | after | | 0.891 | 0.208 | 0.483 | 1.298 |
| 1 | after | | 0.893 | 0.225 | 0.452 | 1.335 |
| 0 | before | | 0.771 | 0.240 | 0.300 | 1.241 |
| 1 | before | | 1.096 | 0.274 | 0.558 | 1.634 |
| Treatment * In prox. area | | | | | | |
| H | | 0 | -0.230 | 0.147 | -0.518 | 0.057 |
| H+ | | 0 | -0.278 | 0.134 | -0.542 | -0.015 |
| H | | 1 | 0.277 | 0.162 | -0.042 | 0.595 |
| H+ | | 1 | -0.044 | 0.139 | -0.316 | 0.229 |

| | | | | | | |
|---|---|---|---|---|---|---|
| In prox. area \| Batch | | | | | | |
| 0 | 1 | | -0.136 | 0.135 | -0.400 | 0.128 |
| 1 | 1 | | 0.087 | 0.141 | -0.189 | 0.363 |
| 0 | 2 | | -0.372 | 0.149 | -0.664 | -0.081 |
| 1 | 2 | | 0.146 | 0.162 | -0.171 | 0.463 |
| Conditioning time \| Batch | | | | | | |
| after | 1 | | -0.365 | 0.151 | -0.660 | -0.069 |
| before | 1 | | 0.315 | 0.152 | 0.018 | 0.613 |
| after | 2 | | -1.117 | 0.156 | -1.424 | -0.811 |
| before | 2 | | 0.891 | 0.187 | 0.526 | 1.257 |
| HumanID | | | | | | |
| AH | | | 0.154 | 0.139 | -0.119 | 0.427 |
| AV | | | -0.292 | 0.143 | -0.571 | -0.012 |
| **-VocPC3** | | | | | | |
| Treatment * In prox. area | | | | | | |
| H | 0 | after | -0.022 | 0.096 | -0.211 | 0.166 |
| H+ | 0 | after | -0.249 | 0.090 | -0.426 | -0.073 |
| H | 1 | after | -0.147 | 0.109 | -0.361 | 0.067 |
| H+ | 1 | after | -0.347 | 0.100 | -0.543 | -0.150 |
| H | 0 | before | -0.322 | 0.116 | -0.551 | -0.094 |
| H+ | 0 | before | -0.356 | 0.101 | -0.554 | -0.157 |
| H | 1 | before | -0.057 | 0.150 | -0.350 | 0.236 |
| H+ | 1 | before | -0.427 | 0.105 | -0.634 | -0.221 |
| Conditioning time \| Batch | | | | | | |
| after | 1 | | -0.483 | 0.092 | -0.664 | -0.303 |
| before | 1 | | -0.818 | 0.094 | -1.001 | -0.634 |
| after | 2 | | 0.100 | 0.095 | -0.086 | 0.287 |
| before | 2 | | 0.236 | 0.122 | -0.002 | 0.475 |
| HumanID | | | | | | |
| AH | | | -0.190 | 0.081 | -0.349 | -0.032 |
| AV | | | -0.292 | 0.084 | -0.456 | -0.127 |
| **Model #4 : Behavioural response during sessions of the conditioning** | | | | | | |
| **CondPC1** | | | | | | |
| Trial number | | | | | | |
| - | | | 0.2 | 0.03 | 0.15 | 0.25 |
| Treatment | | | | | | |
| H | | | -0.080 | 0.240 | -0.562 | 0.402 |
| H+ | | | 0.064 | 0.242 | -0.422 | 0.550 |
| HumanID | | | | | | |
| AH | | | -0.206 | 0.243 | -0.693 | 0.281 |
| AV | | | 0.190 | 0.245 | -0.300 | 0.681 |
| Batch | | | | | | |
| 1 | | | 0.658 | 0.243 | 0.171 | 1.145 |
| 2 | | | -0.674 | 0.245 | -1.165 | -0.183 |
| **CondPC2** | | | | | | |

| | | | | | |
|---|---|---|---|---|---|
| Trial number | | | | | |
| - | | -0.17 | 0.02 | -0.22 | -0.12 |
| Treatment | | | | | |
| H | | 0.385 | 0.152 | 0.080 | 0.690 |
| H+ | | -0.363 | 0.154 | -0.672 | -0.054 |
| HumanID | | | | | |
| AH | | 0.061 | 0.153 | -0.246 | 0.368 |
| AV | | -0.039 | 0.156 | -0.352 | 0.273 |
| Batch | | | | | |
| 1 | | -0.312 | 0.153 | -0.619 | -0.004 |
| 2 | | 0.333 | 0.156 | 0.021 | 0.646 |
| **CondPC3** | | | | | |
| Trial number | | | | | |
| - | | -0.001 | 0.017 | -0.035 | 0.032 |
| Treatment | | | | | |
| H | | -0.176 | 0.103 | -0.383 | 0.031 |
| H+ | | 0.188 | 0.105 | -0.022 | 0.398 |
| HumanID | | | | | |
| AH | | -0.043 | 0.104 | -0.252 | 0.165 |
| AV | | 0.055 | 0.106 | -0.157 | 0.267 |
| Batch | | | | | |
| 1 | | 0.010 | 0.104 | -0.198 | 0.219 |
| 2 | | 0.002 | 0.106 | -0.210 | 0.214 |
| **Occurence of missed contact from human** | | | | | |
| Treatment | | | | | |
| H | - | -0.110 | 0.188 | -0.479 | 0.259 |
| H+ | - | -0.922 | 0.206 | -1.325 | -0.519 |
| Trial \| Batch | | | | | |
| 1 | 2 to 11 | -0.303 | 0.062 | -0.425 | -0.182 |
| 2 | 2 to 11 | -0.127 | 0.059 | -0.243 | -0.010 |
| **Model #5 : Vocal response during session of the conditioning Trial number * Treatment * In prox. area** | | | | | |
| **Call duration (s) (log)** | | | | | |
| Treatment * In prox. area | | | | | |
| H | 0 | -1.263 | 0.038 | -1.338 | -1.189 |
| H+ | 0 | -1.385 | 0.037 | -1.457 | -1.313 |
| H | 1 | -1.621 | 0.050 | -1.719 | -1.522 |
| H+ | 1 | -1.567 | 0.044 | -1.654 | -1.481 |
| Trial number \| Batch | | | | | |
| 1 | | -0.053 | 0.009 | -0.070 | -0.036 |
| 2 | | -0.025 | 0.009 | -0.042 | -0.008 |
| Treatment \| Batch | | | | | |
| H | 1 | -1.621 | 0.053 | -1.724 | -1.517 |
| H+ | 1 | -1.523 | 0.055 | -1.631 | -1.416 |
| H | 2 | -1.264 | 0.060 | -1.381 | -1.146 |
| H+ | 2 | -1.429 | 0.052 | -1.532 | -1.326 |

| | | | | | |
|---|---|---|---|---|---|
| HumanID | | | | | |
| AH | | -0.046 | 0.009 | -0.063 | -0.029 |
| AV | | -0.032 | 0.009 | -0.050 | -0.015 |
| **-VocPC1** | | | | | |
| Trial number * In prox. area | | | | | |
| 0 | | 0.058 | 0.018 | 0.023 | 0.094 |
| 1 | | 0.109 | 0.031 | 0.048 | 0.169 |
| Treatment \| Batch | | | | | |
| H | 1 | -0.053 | 0.219 | -0.481 | 0.375 |
| H+ | 1 | -0.670 | 0.227 | -1.115 | -0.225 |
| H | 2 | -2.570 | 0.248 | -3.056 | -2.084 |
| H+ | 2 | -2.033 | 0.218 | -2.460 | -1.606 |
| In prox. area \| Batch | | | | | |
| 0 | 1 | -0.507 | 0.153 | -0.807 | -0.207 |
| 1 | 1 | -0.216 | 0.183 | -0.575 | 0.144 |
| 0 | 2 | -2.210 | 0.155 | -2.513 | -1.906 |
| 1 | 2 | -2.393 | 0.213 | -2.811 | -1.975 |
| HumanID | | | | | |
| AH | | 0.091 | 0.027 | 0.037 | 0.144 |
| AV | | 0.076 | 0.028 | 0.021 | 0.132 |
| **VocPC2** | | | | | |
| Treatment * In prox. area | | | | | |
| H+ | 1 | -1.372 | 0.181 | -1.727 | -1.018 |
| H | 0 | 0.124 | 0.107 | -0.085 | 0.333 |
| H+ | 0 | 0.220 | 0.104 | 0.016 | 0.425 |
| H | 1 | 0.525 | 0.130 | 0.269 | 0.780 |
| H+ | 1 | 0.340 | 0.119 | 0.108 | 0.572 |
| Trial number * In prox. area | | | | | |
| 0 | | -0.091 | 0.010 | -0.110 | -0.072 |
| 1 | | -0.054 | 0.017 | -0.088 | -0.021 |
| Trial number \| Batch | | | | | |
| 1 | | -0.045 | 0.015 | -0.074 | -0.015 |
| 2 | | -0.100 | 0.015 | -0.130 | -0.071 |
| In prox. area * HumanID | | | | | |
| 0 | AH | 0.317 | 0.106 | 0.110 | 0.524 |
| 1 | AH | 0.402 | 0.123 | 0.161 | 0.643 |
| 0 | AV | 0.027 | 0.107 | -0.182 | 0.236 |
| 1 | AV | 0.462 | 0.128 | 0.212 | 0.712 |
| **VocPC3** | | | | | |
| Trial number * HumanID | | | | | |
| AH | | -0.048 | 0.011 | -0.070 | -0.026 |
| AV | | -0.007 | 0.012 | -0.031 | 0.016 |
| Treatment | | | | | |

| | | | | |
|---|---|---|---|---|
| H | 0.193 | 0.082 | 0.033 | 0.353 |
| H+ | 0.121 | 0.076 | -0.029 | 0.270 |
| In prox. area | | | | |
| 0 | 0.205 | 0.052 | 0.103 | 0.308 |
| 1 | 0.108 | 0.069 | -0.026 | 0.243 |
| Batch | | | | |
| 1 | -0.181 | 0.077 | -0.333 | -0.030 |
| 2 | 0.495 | 0.083 | 0.332 | 0.659 |

Table S4 : Table of raw values of acoustic parameters in each comparison group of interest.

| **Comparison of Isolation Reunion (Isolation/Reunion test : static and silent human)** | | | | | **Effect of proximity during reunion (Isolation/Reunion test, static and silent human)** | | | | | **Effect of proximity sessions of additional positive contacts (conditioning, interacting human)** | | | | |
|---|---|---|---|---|---|---|---|---|---|---|---|---|---|---|
| **Conditioning time** | **Treatment** | **Phase** | | **Ncalls** | **Conditioning time** | **Treatment** | **In prox. area** | | **Ncalls** | **Time in conditioning** | **Treatment** | **In prox. area** | | **Ncalls** |
| **Number of calls per group** | | | | | | | | | | | | | | |
| after | H | iso | | 1204 | after | H | 0 | | 1482 | early | H | 0 | | 1240 |
| | | reuH | | 1976 | | | 1 | | 484 | | | 1 | | 164 |
| | H+ | iso | | 1015 | | H+ | 0 | | 1531 | | H+ | 0 | | 1692 |
| | | reuH | | 2163 | | | 1 | | 568 | | | 1 | | 222 |
| before | H | iso | | 842 | before | H | 0 | | 432 | late | H | 0 | | 779 |
| | | reuH | | 662 | | | 1 | | 226 | | | 1 | | 77 |
| | H+ | iso | | 630 | | H+ | 0 | | 609 | | H+ | 0 | | 865 |
| | | reuH | | 1251 | | | 1 | | 706 | | | 1 | | 129 |
| | | | **Mean of parameter** | **Sd of parameter** | | | | **Mean of parameter** | **Sd of parameter** | | | | **Mean of parameter** | **Sd of parameter** |
| **Mean Dominant Frequency (kHz)** | | | | | | | | | | | | | | |
| after | H | iso | 0.304 | 0.071 | after | H | 0 | 0.320 | 0.087 | early | H | 0 | 0.314 | 0.039 |
| | | reuH | 0.324 | 0.092 | | | 1 | 0.337 | 0.105 | | | 1 | 0.327 | 0.036 |
| | H+ | iso | 0.302 | 0.064 | | H+ | 0 | 0.314 | 0.086 | | H+ | 0 | 0.322 | 0.041 |
| | | reuH | 0.320 | 0.093 | | | 1 | 0.335 | 0.103 | | | 1 | 0.329 | 0.040 |
| before | H | iso | 0.322 | 0.065 | before | H | 0 | 0.334 | 0.080 | late | H | 0 | 0.303 | 0.037 |
| | | reuH | 0.350 | 0.098 | | | 1 | 0.381 | 0.120 | | | 1 | 0.324 | 0.039 |
| | H+ | iso | 0.342 | 0.073 | | H+ | 0 | 0.337 | 0.068 | | H+ | 0 | 0.299 | 0.035 |
| | | reuH | 0.343 | 0.065 | | | 1 | 0.348 | 0.060 | | | 1 | 0.331 | 0.057 |
| **Min F peak (kHz)** | | | | | | | | | | | | | | |
| after | H | iso | 0.286 | 0.049 | after | H | 0 | 0.296 | 0.046 | early | H | 0 | 0.309 | 0.052 |
| | | reuH | 0.299 | 0.050 | | | 1 | 0.308 | 0.061 | | | 1 | 0.325 | 0.050 |
| | H+ | iso | 0.288 | 0.053 | | H+ | 0 | 0.288 | 0.057 | | H+ | 0 | 0.322 | 0.052 |
| | | reuH | 0.293 | 0.058 | | | 1 | 0.306 | 0.058 | | | 1 | 0.324 | 0.050 |
| before | H | iso | 0.315 | 0.051 | before | H | 0 | 0.327 | 0.063 | late | H | 0 | 0.296 | 0.045 |
| | | reuH | 0.333 | 0.062 | | | 1 | 0.345 | 0.059 | | | 1 | 0.316 | 0.040 |
| | H+ | iso | 0.334 | 0.052 | | H+ | 0 | 0.330 | 0.052 | | H+ | 0 | 0.296 | 0.045 |
| | | reuH | 0.336 | 0.049 | | | 1 | 0.342 | 0.043 | | | 1 | 0.318 | 0.047 |

| | | | | | | | | | | | | | | |
|---|---|---|---|---|---|---|---|---|---|---|---|---|---|---|
| **Mac F peak (kHz)** | | | | | | | | | | | | | | |
| after | H | iso | 0.931 | 1.071 | after | H | 0 | 1.151 | 1.342 | early | H | 0 | 0.731 | 0.892 |
| | | reuH | 1.177 | 1.383 | | | 1 | 1.261 | 1.499 | | | 1 | 0.979 | 1.070 |
| | H+ | iso | 0.821 | 1.068 | | H+ | 0 | 1.045 | 1.284 | | H+ | 0 | 0.677 | 0.750 |
| | | reuH | 1.058 | 1.282 | | | 1 | 1.054 | 1.282 | | | 1 | 0.827 | 0.916 |
| before | H | iso | 0.969 | 1.233 | before | H | 0 | 0.911 | 1.161 | late | H | 0 | 0.804 | 0.975 |
| | | reuH | 1.080 | 1.346 | | | 1 | 1.419 | 1.600 | | | 1 | 1.013 | 1.121 |
| | H+ | iso | 0.794 | 1.005 | | H+ | 0 | 0.874 | 1.163 | | H+ | 0 | 0.788 | 0.886 |
| | | reuH | 0.844 | 1.136 | | | 1 | 0.786 | 1.060 | | | 1 | 1.040 | 1.070 |
| **Mode (Hz)** | | | | | | | | | | | | | | |
| after | H | iso | 291.278 | 59.014 | after | H | 0 | 302.674 | 51.629 | early | H | 0 | 322.410 | 49.159 |
| | | reuH | 305.321 | 52.397 | | | 1 | 313.777 | 54.076 | | | 1 | 340.177 | 42.013 |
| | H+ | iso | 292.890 | 45.872 | | H+ | 0 | 295.295 | 59.358 | | H+ | 0 | 332.720 | 49.270 |
| | | reuH | 301.645 | 67.420 | | | 1 | 317.938 | 84.665 | | | 1 | 336.006 | 50.457 |
| before | H | iso | 321.339 | 50.148 | before | H | 0 | 335.165 | 72.839 | late | H | 0 | 303.054 | 45.965 |
| | | reuH | 346.629 | 92.602 | | | 1 | 368.842 | 119.373 | | | 1 | 322.763 | 40.934 |
| | H+ | iso | 340.806 | 53.094 | | H+ | 0 | 335.019 | 47.114 | | H+ | 0 | 303.363 | 45.433 |
| | | reuH | 342.822 | 49.077 | | | 1 | 350.652 | 47.117 | | | 1 | 326.297 | 45.992 |
| **Mean (Hz)** | | | | | | | | | | | | | | |
| after | H | iso | 1817.653 | 385.617 | after | H | 0 | 1885.087 | 426.846 | early | H | 0 | 1494.859 | 294.318 |
| | | reuH | 1879.520 | 423.215 | | | 1 | 1868.693 | 410.882 | | | 1 | 1628.225 | 304.585 |
| | H+ | iso | 1842.219 | 428.577 | | H+ | 0 | 1887.656 | 457.439 | | H+ | 0 | 1443.195 | 263.108 |
| | | reuH | 1878.859 | 453.032 | | | 1 | 1837.232 | 442.185 | | | 1 | 1472.662 | 285.439 |
| before | H | iso | 1769.750 | 442.237 | before | H | 0 | 1811.102 | 443.617 | late | H | 0 | 1524.356 | 294.879 |
| | | reuH | 1822.634 | 433.496 | | | 1 | 1851.117 | 414.251 | | | 1 | 1581.281 | 341.346 |
| | H+ | iso | 1687.113 | 390.208 | | H+ | 0 | 1812.031 | 445.991 | | H+ | 0 | 1507.883 | 269.191 |
| | | reuH | 1786.125 | 457.160 | | | 1 | 1736.176 | 462.373 | | | 1 | 1563.601 | 289.547 |
| **Q50 (Hz)** | | | | | | | | | | | | | | |
| after | H | iso | 750.232 | 484.174 | after | H | 0 | 847.308 | 569.306 | early | H | 0 | 493.711 | 212.211 |
| | | reuH | 842.007 | 554.088 | | | 1 | 831.133 | 505.754 | | | 1 | 567.891 | 232.048 |
| | H+ | iso | 778.814 | 545.853 | | H+ | 0 | 865.811 | 614.602 | | H+ | 0 | 473.187 | 152.404 |
| | | reuH | 858.754 | 600.966 | | | 1 | 813.264 | 558.239 | | | 1 | 494.802 | 159.335 |
| before | H | iso | 751.721 | 524.095 | before | H | 0 | 766.585 | 539.701 | late | H | 0 | 500.546 | 223.284 |
| | | reuH | 803.559 | 544.105 | | | 1 | 881.844 | 547.431 | | | 1 | 581.590 | 319.693 |
| | H+ | iso | 665.355 | 454.203 | | H+ | 0 | 755.510 | 511.427 | | H+ | 0 | 496.292 | 213.039 |
| | | reuH | 742.369 | 506.089 | | | 1 | 709.311 | 487.096 | | | 1 | 555.141 | 238.753 |
| **Q25 (Hz)** | | | | | | | | | | | | | | |
| after | H | iso | 302.742 | 69.646 | after | H | 0 | 330.530 | 103.016 | early | H | 0 | 287.444 | 46.594 |
| | | reuH | 330.573 | 101.286 | | | 1 | 331.723 | 96.435 | | | 1 | 309.890 | 50.641 |
| | H+ | iso | 304.540 | 75.870 | | H+ | 0 | 324.614 | 99.783 | | H+ | 0 | 290.177 | 45.347 |
| | | reuH | 328.360 | 100.583 | | | 1 | 333.044 | 95.537 | | | 1 | 301.554 | 51.488 |
| before | H | iso | 317.037 | 82.577 | before | H | 0 | 328.490 | 95.369 | late | H | 0 | 283.892 | 41.855 |
| | | reuH | 345.858 | 104.483 | | | 1 | 380.512 | 112.884 | | | 1 | 320.480 | 64.909 |
| | H+ | iso | 324.410 | 81.132 | | H+ | 0 | 333.849 | 77.673 | | H+ | 0 | 281.491 | 39.744 |

| | | | | | | | | | | | | | | |
|---|---|---|---|---|---|---|---|---|---|---|---|---|---|---|
| | | reuH | 335.275 | 74.787 | | | 1 | 333.792 | 70.761 | | | 1 | 307.556 | 57.016 |
| **Q75 (Hz)** | | | | | | | | | | | | | | |
| after | H | iso | 2724.508 | 901.531 | after | H | 0 | 2855.251 | 962.026 | early | H | 0 | 1907.359 | 821.008 |
| | | reuH | 2838.172 | 964.096 | | | 1 | 2800.454 | 966.843 | | | 1 | 2271.884 | 873.368 |
| | H+ | iso | 2797.501 | 989.550 | | H+ | 0 | 2871.378 | 1036.874 | | H+ | 0 | 1795.251 | 756.385 |
| | | reuH | 2841.678 | 1032.082 | | | 1 | 2731.682 | 1032.696 | | | 1 | 1886.704 | 797.162 |
| before | H | iso | 2667.213 | 1043.411 | before | H | 0 | 2718.417 | 1054.461 | late | H | 0 | 2005.637 | 806.336 |
| | | reuH | 2716.523 | 1026.942 | | | 1 | 2724.730 | 979.535 | | | 1 | 2132.724 | 873.945 |
| | H+ | iso | 2483.126 | 934.606 | | H+ | 0 | 2770.862 | 1082.494 | | H+ | 0 | 2019.569 | 718.732 |
| | | reuH | 2669.632 | 1139.513 | | | 1 | 2509.778 | 1182.463 | | | 1 | 2167.271 | 749.616 |
| **Centroid (Hz)** | | | | | | | | | | | | | | |
| after | H | iso | 1817.653 | 385.617 | after | H | 0 | 1885.087 | 426.846 | early | H | 0 | 1494.859 | 294.318 |
| | | reuH | 1879.520 | 423.215 | | | 1 | 1868.693 | 410.882 | | | 1 | 1628.225 | 304.585 |
| | H+ | iso | 1842.219 | 428.577 | | H+ | 0 | 1887.656 | 457.439 | | H+ | 0 | 1443.195 | 263.108 |
| | | reuH | 1878.859 | 453.032 | | | 1 | 1837.232 | 442.185 | | | 1 | 1472.662 | 285.439 |
| before | H | iso | 1769.750 | 442.237 | before | H | 0 | 1811.102 | 443.617 | late | H | 0 | 1524.356 | 294.879 |
| | | reuH | 1822.634 | 433.496 | | | 1 | 1851.117 | 414.251 | | | 1 | 1581.281 | 341.346 |
| | H+ | iso | 1687.113 | 390.208 | | H+ | 0 | 1812.031 | 445.991 | | H+ | 0 | 1507.883 | 269.191 |
| | | reuH | 1786.125 | 457.160 | | | 1 | 1736.176 | 462.373 | | | 1 | 1563.601 | 289.547 |
| **Sd (Hz)** | | | | | | | | | | | | | | |
| after | H | iso | 2134.495 | 176.184 | after | H | 0 | 2145.795 | 174.256 | early | H | 0 | 1990.877 | 174.232 |
| | | reuH | 2144.085 | 175.255 | | | 1 | 2141.372 | 177.553 | | | 1 | 2047.922 | 175.263 |
| | H+ | iso | 2153.259 | 199.831 | | H+ | 0 | 2147.761 | 186.968 | | H+ | 0 | 1942.072 | 169.828 |
| | | reuH | 2140.791 | 187.566 | | | 1 | 2120.019 | 192.350 | | | 1 | 1940.150 | 184.449 |
| before | H | iso | 2069.712 | 201.094 | before | H | 0 | 2111.660 | 193.862 | late | H | 0 | 2007.860 | 175.699 |
| | | reuH | 2096.807 | 194.727 | | | 1 | 2070.054 | 195.140 | | | 1 | 1992.677 | 170.148 |
| | H+ | iso | 2022.945 | 173.442 | | H+ | 0 | 2106.114 | 205.720 | | H+ | 0 | 1974.469 | 164.606 |
| | | reuH | 2095.007 | 211.699 | | | 1 | 2071.601 | 217.408 | | | 1 | 1964.886 | 163.211 |
| **IQR (Hz)** | | | | | | | | | | | | | | |
| after | H | iso | 2421.766 | 863.329 | after | H | 0 | 2524.720 | 902.648 | early | H | 0 | 1619.915 | 797.974 |
| | | reuH | 2507.599 | 906.090 | | | 1 | 2468.731 | 912.873 | | | 1 | 1961.994 | 849.624 |
| | H+ | iso | 2492.962 | 947.609 | | H+ | 0 | 2546.763 | 981.693 | | H+ | 0 | 1505.074 | 739.851 |

| | | | | | | | | | | | | | | |
|---|---|---|---|---|---|---|---|---|---|---|---|---|---|---|
| | | reuH | 2513.318 | 978.741 | | | 1 | 2398.638 | 986.570 | | | 1 | 1585.150 | 779.768 |
| before | H | iso | 2350.176 | 993.024 | before | H | 0 | 2389.927 | 1001.070 | late | H | 0 | 1721.745 | 785.012 |
| | | reuH | 2370.665 | 976.677 | | | 1 | 2344.219 | 935.061 | | | 1 | 1812.245 | 828.439 |
| | H+ | iso | 2158.716 | 893.869 | | H+ | 0 | 2437.013 | 1036.722 | | H+ | 0 | 1738.078 | 702.471 |
| | | reuH | 2334.358 | 1094.079 | | | 1 | 2175.986 | 1135.706 | | | 1 | 1859.715 | 720.425 |
| **Sfm** | | | | | | | | | | | | | | |
| after | H | iso | 0.545 | 0.112 | after | H | 0 | 0.565 | 0.121 | early | H | 0 | 0.447 | 0.091 |
| | | reuH | 0.563 | 0.120 | | | 1 | 0.560 | 0.117 | | | 1 | 0.485 | 0.095 |
| | H+ | iso | 0.551 | 0.121 | | H+ | 0 | 0.564 | 0.129 | | H+ | 0 | 0.432 | 0.083 |
| | | reuH | 0.562 | 0.129 | | | 1 | 0.549 | 0.127 | | | 1 | 0.437 | 0.089 |
| before | H | iso | 0.531 | 0.129 | before | H | 0 | 0.541 | 0.131 | late | H | 0 | 0.454 | 0.088 |
| | | reuH | 0.544 | 0.128 | | | 1 | 0.552 | 0.123 | | | 1 | 0.468 | 0.104 |
| | H+ | iso | 0.506 | 0.115 | | H+ | 0 | 0.538 | 0.130 | | H+ | 0 | 0.454 | 0.085 |
| | | reuH | 0.530 | 0.134 | | | 1 | 0.515 | 0.136 | | | 1 | 0.467 | 0.090 |
| **Sh** | | | | | | | | | | | | | | |
| after | H | iso | 0.810 | 0.064 | after | H | 0 | 0.820 | 0.068 | early | H | 0 | 0.757 | 0.058 |
| | | reuH | 0.820 | 0.067 | | | 1 | 0.821 | 0.065 | | | 1 | 0.781 | 0.056 |
| | H+ | iso | 0.810 | 0.068 | | H+ | 0 | 0.817 | 0.072 | | H+ | 0 | 0.751 | 0.052 |
| | | reuH | 0.816 | 0.072 | | | 1 | 0.811 | 0.070 | | | 1 | 0.756 | 0.054 |
| before | H | iso | 0.802 | 0.071 | before | H | 0 | 0.804 | 0.073 | late | H | 0 | 0.761 | 0.056 |
| | | reuH | 0.810 | 0.071 | | | 1 | 0.824 | 0.064 | | | 1 | 0.775 | 0.062 |
| | H+ | iso | 0.788 | 0.064 | | H+ | 0 | 0.803 | 0.071 | | H+ | 0 | 0.762 | 0.055 |
| | | reuH | 0.797 | 0.074 | | | 1 | 0.789 | 0.075 | | | 1 | 0.775 | 0.056 |
| **Entropy H** | | | | | | | | | | | | | | |
| after | H | iso | 0.626 | 0.049 | after | H | 0 | 0.625 | 0.049 | early | H | 0 | 0.595 | 0.044 |
| | | reuH | 0.624 | 0.049 | | | 1 | 0.624 | 0.049 | | | 1 | 0.607 | 0.044 |
| | H+ | iso | 0.624 | 0.051 | | H+ | 0 | 0.622 | 0.054 | | H+ | 0 | 0.589 | 0.040 |
| | | reuH | 0.621 | 0.054 | | | 1 | 0.614 | 0.053 | | | 1 | 0.592 | 0.042 |
| before | H | iso | 0.613 | 0.050 | before | H | 0 | 0.613 | 0.053 | late | H | 0 | 0.595 | 0.042 |
| | | reuH | 0.614 | 0.051 | | | 1 | 0.618 | 0.047 | | | 1 | 0.604 | 0.046 |
| | H+ | iso | 0.602 | 0.047 | | H+ | 0 | 0.603 | 0.053 | | H+ | 0 | 0.595 | 0.042 |
| | | reuH | 0.597 | 0.056 | | | 1 | 0.589 | 0.057 | | | 1 | 0.602 | 0.044 |
| **Skewness** | | | | | | | | | | | | | | |
| after | H | iso | 4.525 | 0.678 | after | H | 0 | 4.485 | 0.663 | early | H | 0 | 4.439 | 0.560 |
| | | reuH | 4.442 | 0.690 | | | 1 | 4.302 | 0.754 | | | 1 | 4.405 | 0.562 |
| | H+ | iso | 4.637 | 0.598 | | H+ | 0 | 4.592 | 0.669 | | H+ | 0 | 4.467 | 0.535 |
| | | reuH | 4.557 | 0.689 | | | 1 | 4.516 | 0.721 | | | 1 | 4.417 | 0.578 |
| before | H | iso | 4.408 | 0.556 | before | H | 0 | 4.473 | 0.578 | late | H | 0 | 4.587 | 0.501 |
| | | reuH | 4.320 | 0.743 | | | 1 | 4.017 | 0.915 | | | 1 | 4.443 | 0.638 |
| | H+ | iso | 4.527 | 0.528 | | H+ | 0 | 4.510 | 0.533 | | H+ | 0 | 4.630 | 0.504 |
| | | reuH | 4.549 | 0.545 | | | 1 | 4.579 | 0.543 | | | 1 | 4.488 | 0.594 |
| **Kurtosis** | | | | | | | | | | | | | | |
| after | H | iso | 24.851 | 6.106 | after | H | 0 | 24.571 | 5.721 | early | H | 0 | 23.585 | 5.285 |
| | | reuH | 24.187 | 5.944 | | | 1 | 22.948 | 6.454 | | | 1 | 23.511 | 5.443 |
| | H+ | iso | 25.901 | 5.610 | | H+ | 0 | 25.663 | 5.917 | | H+ | 0 | 23.892 | 5.198 |
| | | reuH | 25.329 | 6.093 | | | 1 | 24.865 | 6.429 | | | 1 | 23.557 | 5.393 |

| | | | | | | | | | | | | | | |
|---|---|---|---|---|---|---|---|---|---|---|---|---|---|---|
| before | H | iso | 23.361 | 5.093 | before | H | 0 | 24.071 | 5.215 | late | H | 0 | 25.056 | 4.700 |
| | | reuH | 22.971 | 6.224 | | | 1 | 20.784 | 7.356 | | | 1 | 24.180 | 5.847 |
| | H+ | iso | 24.464 | 5.004 | | H+ | 0 | 24.275 | 4.880 | | H+ | 0 | 25.495 | 4.883 |
| | | reuH | 24.699 | 5.008 | | | 1 | 25.036 | 5.006 | | | 1 | 24.380 | 5.485 |
| **Call duration (s)** | | | | | | | | | | | | | | |
| after | H | iso | 0.497 | 0.252 | after | H | 0 | 0.366 | 0.193 | early | H | 0 | 0.358 | 0.178 |
| | | reuH | 0.366 | 0.205 | | | 1 | 0.364 | 0.236 | | | 1 | 0.243 | 0.166 |
| | H+ | iso | 0.435 | 0.164 | | H+ | 0 | 0.349 | 0.163 | | H+ | 0 | 0.330 | 0.174 |
| | | reuH | 0.339 | 0.161 | | | 1 | 0.316 | 0.156 | | | 1 | 0.265 | 0.151 |
| before | H | iso | 0.387 | 0.195 | before | H | 0 | 0.326 | 0.177 | late | H | 0 | 0.301 | 0.185 |
| | | reuH | 0.308 | 0.180 | | | 1 | 0.272 | 0.181 | | | 1 | 0.236 | 0.177 |
| | H+ | iso | 0.329 | 0.134 | | H+ | 0 | 0.262 | 0.122 | | H+ | 0 | 0.255 | 0.134 |
| | | reuH | 0.248 | 0.123 | | | 1 | 0.231 | 0.120 | | | 1 | 0.203 | 0.111 |

Table S5: Number of calls of each call type recorded during the session and the number of pigs involved in the count. Taking into account the different statistical variable that needed to be add in the models, and thus the number of calls and pigs needed to have reliable statistical analysis, it was thus decided to use only grunts in this study.

| | | **Before conditioning (Isolation/Reunion test – Reunion with H)** | | **During conditioning (all trials pooled)** | | **After conditioning (Isolation/Reunion test – Reunion with H)** | |
|---|---|---|---|---|---|---|---|
| **Call type** | **Treatment** | **N calls** | **N pigs** | **N calls** | **N pigs** | **N calls** | **N pigs** |
| bark | H | 14 | 5 | 13 | 7 | 6 | 3 |
| grunt | H | 670 | 21 | 3979 | 29 | 1981 | 25 |
| mixed | H | 8 | 1 | 172 | 12 | 157 | 9 |
| scream | H | 0 | 0 | 14 | 4 | 39 | 2 |
| squeal | H | 11 | 2 | 94 | 10 | 66 | 11 |
| bark | H+ | 4 | 2 | 18 | 6 | 1 | 1 |
| grunt | H+ | 1244 | 27 | 5006 | 29 | 2072 | 27 |
| mixed | H+ | 0 | 0 | 142 | 12 | 21 | 3 |
| scream | H+ | 0 | 0 | 7 | 2 | 0 | 0 |
| squeal | H+ | 8 | 2 | 50 | 8 | 25 | 6 |
